# Quantitative omnigenic model discovers interpretable genome-wide associations

**DOI:** 10.1101/2024.02.01.578486

**Authors:** Natália Ružičková, Michal Hledík, Gašper Tkačik

**Affiliations:** Institute of Science and Technology Austria, AT-3400 Klosterneuburg, Austria

**Keywords:** genome-wide association study, gene regulation, networks, yeast, the omnigenic model

## Abstract

As their statistical power grows, genome-wide association studies (GWAS) have identified an increasing number of loci underlying quantitative traits of interest. These loci are scattered throughout the genome and are individually responsible only for small fractions of the total heritable trait variance. The recently proposed omnigenic model provides a conceptual framework to explain these observations by postulating that numerous distant loci contribute to each complex trait via effect propagation through intracellular regulatory networks. We formalize this conceptual framework by proposing the “quantitative omnigenic model” (QOM), a statistical model that combines prior knowledge of the regulatory network topology with genomic data. By applying our model to gene expression traits in yeast, we demonstrate that QOM achieves similar gene expression prediction performance to traditional GWAS with hundreds of times less parameters, while simultaneously extracting candidate causal and quantitative chains of effect propagation through the regulatory network for every individual gene. We estimate the fraction of heritable trait variance in *cis-* and in *trans-*, break the latter down by effect propagation order, assess the *trans-* variance not attributable to transcriptional regulation, and show that QOM correctly accounts for the low-dimensional structure of gene expression covariance. We furthermore demonstrate the relevance of QOM for systems biology, by employing it as a statistical test for the quality of regulatory network reconstructions, and linking it to the propagation of non-transcriptional (including environmental) effects.

**Significance statement:** Genetic variation leads to differences in traits implicated in health and disease. Identifying genetic variants associated with these traits is spearheaded by “genome-wide association studies” (GWAS) – statistically rigorous procedures whose power has grown with the number of genotyped samples. Nevertheless, GWAS have a substantial shortcoming: they are ill-equipped to detect the causal basis and reveal the complex systemic mechanisms of polygenic traits. Even a single genetic change can propagate throughout the entire genetic regulatory network causing a myriad of spurious detections, thereby significantly limiting GWAS usefulness. To this end, we propose a novel statistical approach that incorporates known regulatory network information with the potential to boost interpretability of state-of-the-art genomic analyses while simultaneously extracting systems biology insights.

## Introduction

During the past two decades, genome-wide association studies (GWAS) have provided us with a number of key insights into the genetic basis of complex traits [50]. We now know that typical traits are influenced by many loci of small effect; consequently, our inability to fully account for the heritable variance in a trait (“missing heritability”) mainly originates in the statistical limitations of our surveys [22, 6, 36, 18]. Furthermore, there is a high prevalence of significant GWAS hits in the noncoding DNA, especially in regulatory sequences such as enhancers and promoters [46, 23, 26]. The distribution of hits across the genome has been shown to be surprisingly uniform, rather than concentrated in or near “core genes,” i.e., genes with a clear biological and causal link to the trait of interest [12, 38, 49]. Motivated by these insights, Boyle and colleagues proposed the “omnigenic model” for complex traits [6], in which various regulatory networks – transcriptional, post-translational, signaling, etc. – propagate the effects of distant mutations to the core genes and thus to the trait value [14]. Cellular networks are a prime candidate for such propagation, because they are highly interconnected into small-world and scale-free topologies [4, 3, 2]: a particular variant might change the expression level of a proximal gene (as a *cis*-eQTL), this change would in turn affect the expression level of a downstream gene and so on, until essentially any trait of interest could be affected by many mutations via a limited number of intermediate steps. While these effects may be small individually, they will be numerous, providing a possible explanation for the widespread pleiotropy [11, 45] and polygenicity [34, 5, 40], the bias of effect size distribution towards small values, and the uniformity of hits over the genome [47, 49, 42, 30, 8].

Despite its attractiveness, it is unclear how to recast the omnigenic model, which is conceptual at its core, into a statistical model that can be fit to data, rigorously tested, and quantitatively interpreted. If this were possible, we could not only reliably assess the fraction of heritable variance in *cis-* vs. in *trans-* in aggregate [18], but decompose the *trans-* effects order-by-order and identify major chains of effect propagation through regulatory networks for each individual trait. Success in this enterprise would have an important interpretative value and could significantly constrain the hypothetical causal mechanisms that require dedicated followup experiments. Furthermore, it could uncover the causal basis of genetic architecture of complex traits, with broad implications for the fields of evolutionary biology and population genetics [**Fagny2021, Wittkopp2004**, 51]. While the omnigenic model was inspired mainly by observations in human GWAS, we emphasize that its key idea of genetic effect propagation along intracellular networks to phenotypes of interest should hold more broadly and be unrelated to various idiosyncrasies of human genomics. This provided the motivation and context for our study.

In this paper, we propose one possible mathematical formalization of the omnigenic model that we named the “quantitative omnigenic model” (QOM). To demonstrate its potential, we apply it to a subset of 183 transcription factor (TF) genes that form the core of the yeast genetic regulatory network [25]. The rationale for our focus on the TF genes is twofold: first, effect propagation in the remaining yeast genes should be straightforward, since non-TF genes simply inherit the effects propagated from their regulators immediately upstream; second, transcriptional regulation is an essential process involved in mapping genotypes into complex phenotypes in the omnigenic model. The traits of interest in our study are gene expression levels (GE), a common intermediate phenotype in complex trait GWAS [15, 17, 55, 54, 33], so that our results can be directly compared to the traditional eQTL analyses of the same primary data [1].

Previous work that sought to link biological information with statistical genomics can be sorted into three broad groups. In one approach, regulatory network or pathway information can be used *post hoc*, to interpret GWAS findings [43, 13, 39, 48, 14, 53, 40, 27]– rather than include this information into the model *a priori*, as we do. In an alternative, Bayesian approach, side knowledge about the genetic architecture of a trait can be incorporated into marker effect priors [21, 28], or the structure of latent variables responsible for the trait value [29] – while explanatory, these models do not directly probe effect propagation, which is our aim. Lastly, effect propagation has been probed in the frameworks of mediation analysis [32, 52, 37] or structural equation modeling (SEM) [35, 7, 9, 31]. These frameworks seek to infer both the strength *and* topology of the underlying regulatory network from data. While methodologically interesting, statistical limitations render these approaches intractable for larger networks due to the vast space of possible degenerate regulatory topologies that can fit the data, with a consequent drop in explanatory potential. Our key innovation in QOM is to utilize prior data on the topology of regulatory interactions, while leaving the models free to infer the strength of direct (*cis-*) effects as well as the strength of network interactions (*trans-* effects) given the topology, as we describe below.

## Quantitative Omnigenic Model

The Quantitative Omnigenic Model (QOM) can be specified in terms of a *N ×G* matrix of (standardized) gene expression level traits **Y**, where *N* is the number of individuals and *G* is the number of genes of interest, and a *N × P* matrix of genotypes **X**, where *P* is the number of tracked polymorphisms. Given these two matrices, standard GWAS-type model analogous to Polygenic Risk Score (PRS) can be formulated as a linear regression:

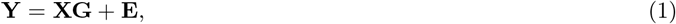

where **G** is a *P × G* matrix of the sought-for additive effects that can be inferred from data, typically by imposing a regularization scheme (sparse / L1, ridge / L2, spike-and-slab, etc.) over effect values to prevent overfitting. **E** is the residuals matrix, usually assumed independent and identically distributed normal, which also includes non-additive genetic, random environmental (“noise”), and other interaction effects.

In contrast to the standard GWAS-type models, QOM requires two further pieces of information, encoded in matrices **D** and **B**. The first is a classification, for each of the *G* genes, whether a given polymorphism could affect that gene’s expression level in *cis* or not. Typically, polymorphisms included in this category would be proximal to the gene, i.e., within some threshold genomic distance (but perhaps also including known distal enhancer regions), capturing the effects on expression due to mutations in the coding, intron, promoter, and proximal enhancer regions. This prior knowledge about possible *cis* effects determines the sparsity pattern of a *P × G* matrix of *direct* effects on gene expression **D**: if a polymorphism *i* cannot affect gene *j* in *cis*, we fix *d*_*ij*_ = 0; the remaining free values of **D** that contain the direct effects of polymorphisms on gene expression levels need to be inferred by QOM.

The second piece of extra information is the putative topology of the regulatory network between the *G* genes of interest. This topology is usually available as a directed graph of *direct* interactions that can be encoded into a *G × G* matrix **B**: we fix *b*_*ij*_ = 0 if gene *i* is not a direct regulator of gene *j*, i.e., if there is no directed interaction from gene *i* to gene *j* in the known regulatory network topology. The remaining free values of **B** need to be inferred by QOM.

Our knowledge of the actual regulatory network topology, encoded into matrix **B**, is probably incomplete, because TF binding measurements uncover only one type of regulatory mechanism; moreover, due to the noisy nature of such experiments the network may contain erroneously reported regulatory links. While this is an important limitation of input data that our approach depends on, we will also develop and deploy various statistical controls to make sure that the network topology we use nevertheless non-trivially contributes to the explanatory power of our studies.

With these definitions in place, QOM can be formulated as a sequence of models schematized in Fig 1A. One starts with a *cis* or the 0^*th*^ *order* QOM, which trivially takes the form of PRS-like linear regression restricted to the *cis* genetic variants for each gene, i.e., **Y** = **XD** + **E**. In the 1^*st*^ *order* QOM, defined as **Y** = **XD** + **XDB** + **E**, the second term propagates the *cis* effects **XD** through one direct step via regulatory matrix **B**. QOM models of order *K* can be constructed in an analogous fashion,

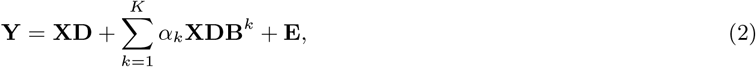

to explicitly model *cis* effect propagation via up to *K* steps through the regulatory network. *α*_*k*_ are scaling parameters which we fix to 1 for reasons explained below, but which in general could be treated as further free parameters. Since matrices **B** and **D** contain unknown effects to be inferred, these problems are manifestly nonlinear even before regularization, but gradient-based large-scale optimization offers a viable numerical method of solution.

**Figure 1:**
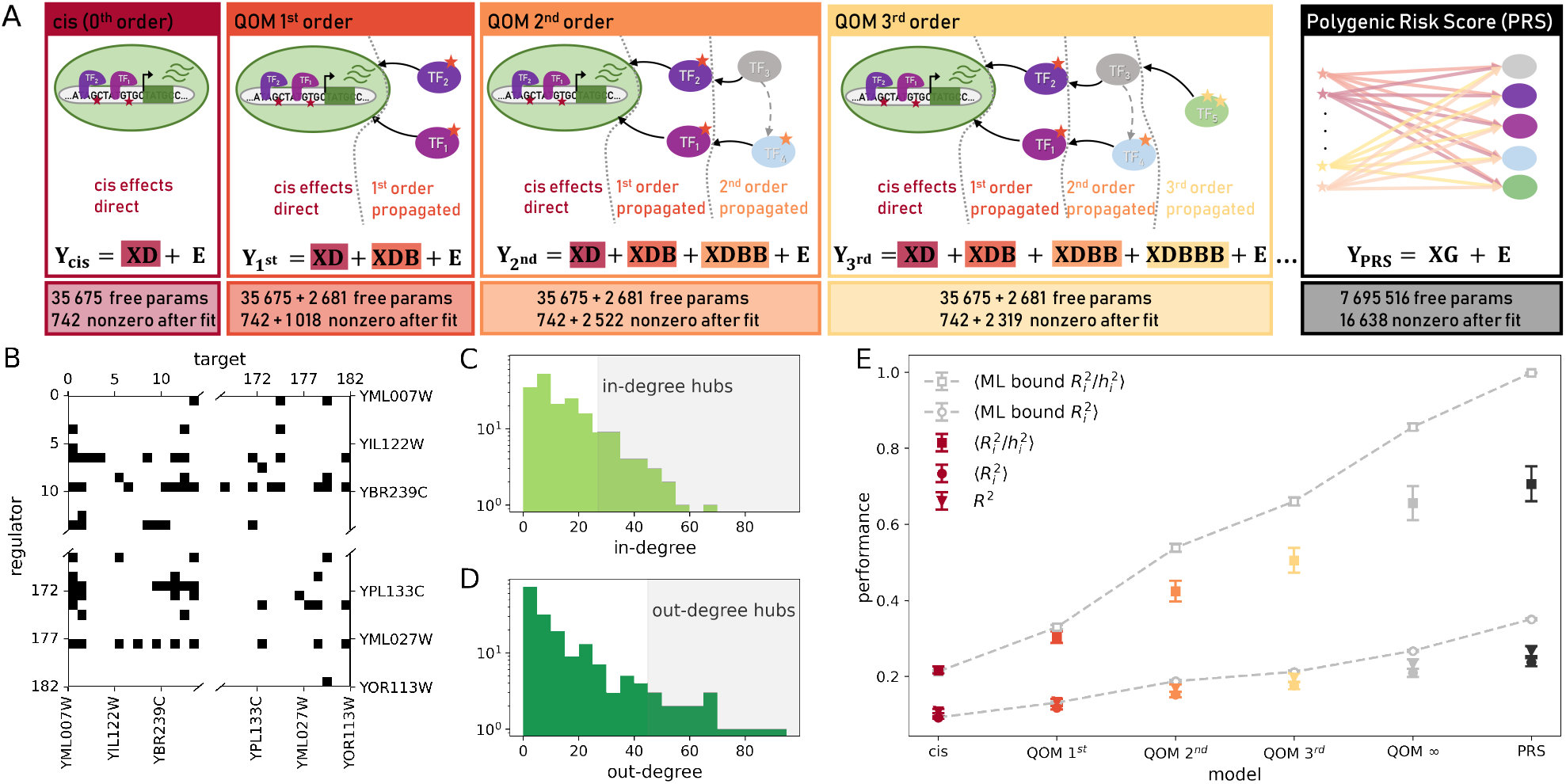
Quantitative omnigenic model (QOM) propagates direct (*cis*) effects on gene expression through a known gene regulatory network order-by-order. **(A)** Color-coded schematics of individual models arranged by increasing order of effect propagation left to right, and the respective numbers of parameters before and after the fit. *cis*: only direct (*cis*) effects, QOM 1^*st*^, 2^*nd*^ and 3^*rd*^ order: direct effects propagated once, twice, three times through the GRN, respectively. The number of free parameters in QOM is the number of direct effects plus the number of links, and remains constant across orders. **Y**: *N* × *G* matrix of gene expression levels (*N* : number of individuals, 1012, *G*: number of TFs, 183), **X**: *N* × *P* genotypes matrix (*P* : number of polymorphisms, 42 052), **D**: *P* × *G* matrix of direct effects on GE levels (*cis*-eQTLs in proximity of genes’ coding sequences), **B**: *G* ×*G* transcriptional regulatory network matrix, structured according to Yeastract+ reported topology [25]. **(B)** A segment of the GRN matrix **B.** Black (8% of matrix entries) = DNA-binding experimental evidence for TF-target interaction, can propagate genetic effects, value to be inferred; white = no evidence for interaction, value fixed to 0. **(C**,**D)** Distribution of in-(out-) degrees of TFs in **B**. In-(out-) degree hubs, highlighted in grey, are defined as TFs with in-(out-) degree larger than 85% (90%) of all genes, equal to 27 (45). **(E)** Measures of model performance for individual models, color as in (A). Total fraction-of-variance in GE explained (*R*^2^, triangle), average per gene fraction-of-variance explained (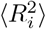, circle), and mean fraction of GE heritability explained (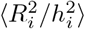,square). The corresponding ML upper bounds (grey) estimate the variance explained by the set of polymorphic sites that contribute to each of the QOM and PRS models (SI Appendix Sec. 3). Model performance errorbars are SD across 50 resamplings of 112 individuals from the evaluation set (SI Appendix Sec. 1). ML errorbars are SD over resamplings of 800 individuals (training set size).

Before proceeding, we make three important remarks. First, the QOM hierarchy can be formally interpreted as a Taylor expansion of the explicit solution for gene expression levels **Y** given by a model with additive contribution of direct and propagated effects (in a form of a SEM):

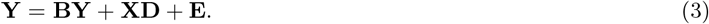

Note that this equation is implicit in **Y**; an explicit linear algebra solution requires computing (I *−* **B**)^*−*1^, whose Taylor expansion generates the QOM sequence of Eq (2), with all *α*_*k*_ = 1 as we assumed. Regulatory networks are, however, not linear; specifically, inputs to a given gene are often processed by saturating (sigmoid) nonlinearities, prompting us to consider generalizations of Eq (3) where the inputs on the right-hand side are passed through some nonlinear function *f* (*·*). Quite generally, such a function could similarly be Taylor-expanded, generating the series of Eq (2) with non-trivial coefficients *α*_*k*_ that likely decay with propagation order *k*. We leave this generic case open for future research.

Second, QOM of higher order (with *α*_*k*_ = 1 which we fix henceforth) do not require any extra free parameters compared to the 1^*st*^ *order* QOM, even though we are modeling multi-step effect propagation and the interaction parameters **B** enter raised to a higher power. Statistically, this provides a solid, albeit unusual, basis for order-by-order model comparison, where the number of parameters is fixed and thus no model complexity penalty is needed. To enable meaningful comparisons, we first fit the *cis* (0^*th*^ *order* QOM) model to extract direct effects by estimating the matrix **D**; for higher order QOM models we do not re-estimate **D** (therefore getting a lower-bound on the models’ performance), but do estimate **B** for each QOM model separately. This enables us to study how effect propagation is captured by models of various orders and compare that to the propagation estimated by the highest-order (*K* = 3) model we consider.

Third, since our goal is not only to identify significant eQTLs but to build interpretable and predictive models, we have formulated PRS and QOM as regression problems rather than as significance tests for individual polymorphic sites. We infer the effects by minimizing the mean square error in predicted gene expression across genes and genotypes, subject to sparse (L1) regularization (SI Appendix Sec. 1). Because gene regulatory matrix **B** couples all the genes, higher-order QOM models cannot be solved sequentially gene-by-gene, but must instead be solved globally. Throughout the manuscript, we will use several metrics for predictive power and model comparison, always evaluated over random withheld (test) subset of genotypes: total fraction-of-variance explained, average per gene fraction-of-variance explained, or average per gene fraction-of-heritable-variance explained. For detailed model specifications, inference, regularization and numerical solution methods, see SI Appendix Sec. 1; we also tested this methodology on synthetic data (SI Appendix Sec. 10, Fig S1).

## Results

### Inferring and testing Quantitative Omnigenic Models

For the main analysis, we applied the QOM framework to the dataset of Albert *et al* [1], who generated 1012 segregants from a *Saccharomyces cerevisiae* yeast cross between laboratory (BY) and wineyard (RM) strain which were genotyped and profiled for gene expression by RNA sequencing (SI Appendix Sec. 2). We restricted the non-zero structure of the direct (*cis*) effects matrix **D** to polymorphisms within confidence intervals of previously identified *cis*-eQTLs [1], and extracted the non-zero structure of the gene regulatory network interaction matrix **B** from the Yeastract+ database [25] based on “DNA-binding evidence” in *Saccharomyces cerevisiae*. This evidence consists mainly of ChIP-derived information and is thus putatively indicative of direct regulation (Fig S2 replicates the key analyses on dataset by Albert et al [1] using Yeastract “expression evidence” instead, SI Appendix Sec. 7.) For the supplementary analysis, we used an entirely different, natural *S. cerevisiae* “pan-population” dataset [**Caudal2024**], also using the DNA-binding evidence network. Key results from this supplementary analysis are shown in Figs S10,S11 and commented on in Discussion.

We restricted both analyses to the sub-network of TFs, which left us with *G* = 183 genes. This gene regulatory network matrix is sparse (Fig 1B), with only *∼* 8% of elements being potentially non-zero; visual inspection suggests that non-zero elements are not distributed randomly but are over-represented in rows that correspond to individual TFs which regulate many downstream TF genes (Fig S3). Distributions of in-degree (Fig 1C) and especially out-degree (Fig 1D) correspondingly have long tails, in line with previous studies in cellular networks [2]; we use these histograms to select in-and out-degree hub genes for subsequent analyses.

We evaluate the performance of QOM and comparison models in Fig 1E; the distributions of inferred effect sizes are reported in Fig S4. Most importantly, we observe a substantial boost in predictive performance by including the ability of *cis* effects to propagate: 0^*th*^ *order* QOM explains 9% of expression variance (average per gene), which rises to 18% for the 3^*rd*^ *order* model. This is approaching the unconstrained PRS (24%) and nearly reaches the upper bound on the performance of QOM models (21%) (denoted by “QOM *∞*”, see SI Appendix Sec. 1 for definition).

We can independently upper bound the fraction of variance in GE explainable for each gene when we do not seek to infer individual effect sizes, using the maximum likelihood (ML) variance estimates (SI Appendix Sec. 3). These upper bounds reveal that the PRS model (unconstrained regression) captures a bit more than two thirds of achievable variance given the sample limitations of the dataset at hand; QOMs capture more than three quarters. We can furthermore ask about the fraction of heritable variance explained, with heritability (*h*^2^) estimates taken from Ref [1]. Here, too, the 3^*rd*^ *order* QOM approaches the QOM bound and is not far from the unconstrained PRS, both of which can account for around 50% of heritable variance in this dataset, on average.

Taken together, these results validate the basic premises of the omnigenic model [6]: propagating direct effects along transcriptional regulatory network is sufficient to achieve ≳ 70% of unconstrained PRS performance in all measures considered, and propagation along only a handful of steps (here, three) nearly saturates QOM performance. This is consistent with the GRN topology: at 3^*rd*^ order, *>* 90% of *trans-*eQTLs can be reached via effect propagation; at 5^*th*^ order, all of them could be reached (Fig S3). QOM performance results are surprisingly good given the severe constraints placed on the models – both in statistical (in terms of free parameters) as well as structural terms (as ultimately *all* effects are either in *cis* to a given gene, or must propagate along a fixed network from other genes’ *cis* effects).

### Effect propagation for individual genes

We next shift focus from aggregate performance measures to effect propagation for individual genes. Figure 2A shows that QOMs are in general better at predicting genes with high heritability. Among genes with above median heritability (*h*^2^ *>* 0.29), the dominant contribution to the increase in predictive power with QOM order comes from genes that are simply not predictable from their *cis* variants and *K* propagation steps, but can suddenly be predicted by propagating effects through one more, (*K* + 1)-th, step.

**Figure 2:**
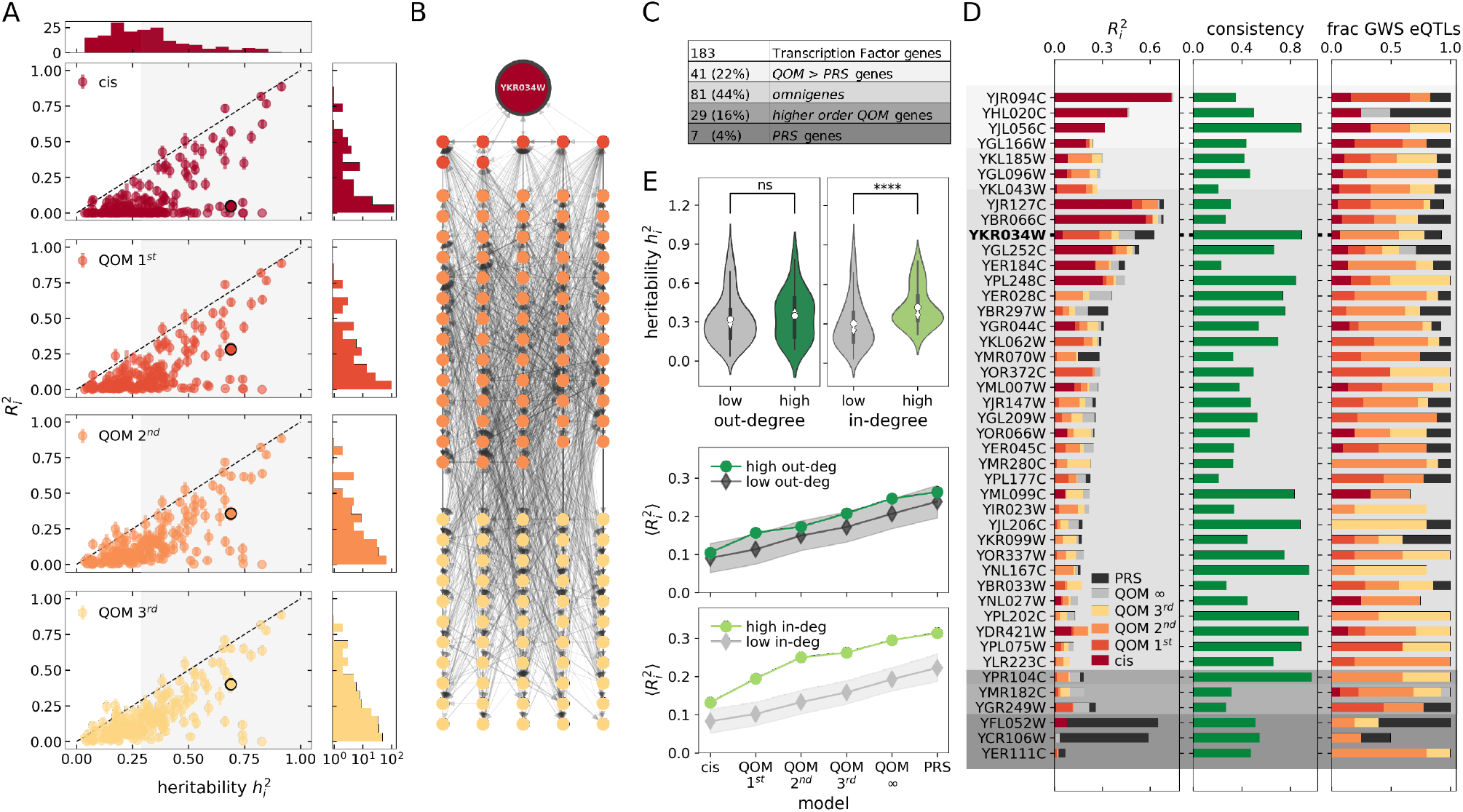
Models including higher orders of effect propagation explain more variance in GE for individual genes. **(A)** Per gene variance explained 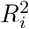 as a function each gene’s heritability 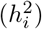, estimated by Albert *et al* [1]. Marginal distributions of 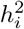 and 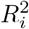 shown on the sides. Example gene `YKR034W` highlighted in black. **(B)** Reconstructed pathways of effect propagation for `YKR034W`, according to 3^*rd*^ *order* QOM. **(C)** Categories of genes. *Omnigenes*: genes where including higher orders of propagation increases 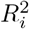 (i.e. QOM 3^*rd*^ *>* QOM 2^*nd*^ *>* QOM 1^*st*^ *> cis*). *Higher order QOM* genes: all lower order models explain less than 60% of QOM ∞ expression variance while QOM ∞ explains at least 80% of PRS variance. *PRS* genes: QOM 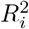 is less than 40% of PRS 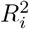. **(D)** Example genes from each category denoted by grey-scale as in (C). Categories can overlap. Per gene 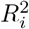, consistency score, and the fraction of genome-wide significant (GWS) eQTLs identified by Ref [1] captured by the models, accounting for LD and eQTL confidence intervals. For clarity, examples with highest 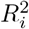 and consistency*>* 0.2 are shown. Averages across all genes for comparison, not plotted: consistency score (0.35), fraction of GWS eQTLs captured by QOM 1^*st*^(0.24); 2^*nd*^(0.61); 3^*rd*^(0.79); ∞ (0.73) and *PRS* (0.91). **(E)** In and out-degree hubs. **Top:** distribution of *h*^2^ for hubs vs non-hub genes. No significant difference between genes with high (⟨*h*^2^⟩ = 0.35) and low (⟨*h*^2^ ⟩= 0.32 ±0.05, 100 resamplings) out-degree (*p >* 0.1, U-test). In-degree hubs have significantly higher *h*^2^ (⟨*h*^2^ ⟩= 0.43) than non-hub genes (⟨*h*^2^ ⟩= 0.30 ±0.04, 100 resamplings; *p <* 10^*−*4^, U-test). White circle signifies the mean, diamond the median. **Middle, Bottom:** 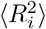 of different models for hubs vs non-hubs. Shaded areas denote standard error of the mean across 100 samples of *H* non-hub genes, where *H* is the number of hub genes (*H* = 29 in- and *H* = 17 out-degree hub genes). Unlike out-degree hubs, in-degree hubs have significantly higher 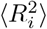 than non-hub genes, the difference being highest for QOM 1^*st*^ − 3^*rd*^.

A case in point is Dal80 (`YKR034W`), a transcriptional regulator involved in yeast nitrogen degradation (Fig 2B). Despite its high heritability (*h*^2^ *≈* 0.7), its expression levels are poorly predictable from *cis-* effects alone; inclusion of one-step contributions from its 7 regulators (out of 8, given the structure of **B**), whose expressions are predicted by their own respective *cis-* variants, combined with the inclusion of 68 two-step (out of 73) and (all) 55 three-step regulators pushes the fraction of heritability explained to *≈* 58%. Overall, the pathways of effect propagation for Dal80 shown in Fig 2B trace back to 276 sites (identified previously as *cis*-eQTLs of some of the 183 TF genes). The majority (176) of those downregulate Dal80 expression, and five are outside the 3SD interval around the mean. Although the interactions propagate through a network of 131 genes, the putative causal sites are *cis-*eQTLs to only 84 genes (and YKR034W itself). The inferred genetic effects of these sites range from *−*0.29 to 0.19. Among top 10 polymorphisms with largest absolute effect size, four are *cis-*eQTLs of “master regulator” out-degree hubs: YMR016C, YNL216W, YNL068C. As all genetic effects are *cis-*eQTLs to TF genes, we looked at the eight genes in whose promoters these top 10 polymorphisms reside. Only one of them is a direct regulator of Dal80, five are 2^*nd*^ order regulators and the remaining two genes enter only at the 3^*rd*^ order.

One can extract and analyze the chains of effect propagation for each of the remaining 182 genes in an analogous fashion, remembering that the inferred interaction strengths *b*_*ij*_ are self-consistently estimated *only once*, jointly for all the expression traits (all genes), when the 3^*rd*^ *order* QOM is fitted to data.

We categorized all 183 TF genes into four categories based on the fraction of their variance explained by QOMs (Fig 2C; SI Appendix Sec. 5). For nearly half of the genes analyzed, QOM outperforms the unrestricted PRS; only for seven genes (“PRS genes”), on the contrary, PRS strongly outperforms QOMs, very likely indicating that their expression, while predictable, cannot be explained by effect propagation through the transcriptional regulatory network. If we exclude “PRS genes” from the analysis, the gap in the average per gene fraction-of-variance explained between QOM 3^*rd*^ *order* and PRS drops by a quarter. Moreover, 51% of the genetic effects identified by the PRS model do not overlap a *cis-*eQTL of any of the TF genes in our GRN. These sites account for nearly 5% of the total expression variance explained by PRS, are responsible for nearly the entire performance gap between PRS and 3^*rd*^ *order* QOM for all genes, and are enriched in non-transcriptional regulatory hotspots [1]. These results provide a rough estimate of fraction of heritable variance not explainable by propagating direct effects along the transcriptional regulatory network – either because the network reconstruction is incomplete; or, as is likely, effects also propagate via other intracellular networks.

For 44% of the analyzed genes, which we refer to as “omnigenes,” the expression level predictability rises with QOMs of increasing order. Despite this rigorous criterion (which excludes genes whose predictability saturates after, e.g., one step of propagation), the high percentage of omnigenes points to the widespread occurrence of effect propagation. A further 16% of genes are “higher order QOM genes” where the QOM bound suggests room for improvement if propagation beyond 3^*rd*^ order were considered.

Figure 2D shows example genes from each category, breaking down expression variance contributions by QOM order (see Fig S5 for a similar analysis of order-by-order variance contributions within a single 3^*rd*^ *order* QOM fit). Looking beyond variance explained, we next wondered how consistent are the pathways of effect propagation identified by different-order QOMs. To this end we defined a consistency score, essentially an average Pearson correlation between effects on a given gene identified by different-order QOMs (SI Appendix Sec. 4). Even though QOMs of higher order can discover *new* pathways that are simply out of reach at lower order – implying that even in the ideal case the expected consistency should be below 1 – we nevertheless frequently observe high consistency scores easily exceeding 0.5. This suggests that strongly overlapping pathways are identified by independently inferred models. Since each such effect propagation pathway for a given gene ultimately traces back to a polymorphism which must be a *cis*-eQTL for one of the TF genes, we asked how strongly these eQTLs themselves overlap with the genome-wide significant (GWS) eQTLs as identified by the standard GWAS analysis by Albert *et al* [1]. The last column of Fig 2D shows that the fraction of GWS eQTLs picked up by QOM is high, on average *∼* 80% for 3^*rd*^ *order* QOM, with the large majority of identified eQTLs in *trans* to a gene of interest. Moreover, there is a substantial overlap between the effect signs of *trans* eQTL identified by a GWAS and by 3^*rd*^ *order* QOM (SI Appendix Sec. 4). QOM therefore not only identifies eQTLs that overlap with standard GWAS results, but tells us how these eQTLs affect their distal targets. While our pathway identification is statistical in nature, it provides specific, testable, causal, and mechanistically-interpretable hypotheses that have been out of reach for alternative approaches.

Is predictability of a given gene expression level correlated with the local topology of the regulatory network around that gene? Figure 2E looks at in-degree and out-degree hubs as defined in Fig 1C,D. While heritability is not significantly different between out-degree hubs and non-hubs, it is significantly higher for in-degree hubs: expression levels of TFs that are regulated by more regulators tend to be more heritable. Correspondingly, ML estimates show that there is significantly more explainable variance for in-degree hubs than for other genes, and we likewise observe that for in-degree hubs QOMs explain significantly more expression variance (and a higher fraction of heritable variance, Fig S6) compared to non-hubs or out-degree hubs.

Taken together, these results paint an interesting interplay between the genetic architecture of expression traits and the regulatory network topology, whereby more incoming regulatory arrows enable more potential loci in *trans* to affect the expression trait. While PRS and QOM both identify the widespread polygenic basis for GE traits, they differ in the inferred degree of polygenicity (Fig S7). In PRS, which relies heavily on sparse regularisation, the number of sites associated with a GE trait is relatively modest (around 10 to 170 eQTLs per gene). In contrast, the 3^*rd*^ *order* QOM suggests that most *cis-*eQTLs affect a large fraction of the other TFs, with most genes having 250-300 eQTLs. A plausible interpretation for this difference is that the prior knowledge of network topology that drastically restricts the structure and number of free parameters allows QOM to identify weaker effects, which the PRS model has to regularize away in order to generalize well. By implication, GE traits may be even more polygenic than the standard GWAS-type analyses suggest.

### QOM as a statistical test for regulatory network topologies

Next, we switch gears to explore the implications of QOMs for systems biology. The field of systems biology is nearly synonymous with the ambition to understand collective biological function emerging from interactions between networked components. Genetic regulatory networks, in particular, are a paradigm for such explorations, with extensive efforts dedicated to reconstructing the topology of transcriptional interactions in model systems such as *Escherichia coli* [44], yeast [25, 10], *Drosophila* [16], and mammals, including human [19, 20]. While these networks are often inferred painstakingly either link by link in dedicated molecular biology experiments, or through large and expensive genomic-scale experimental efforts, to our knowledge there is no systems-level methodology to evaluate the quality and predictive power of such network-level reconstructions.

Because the postulated network topology structures QOM regression, we reasoned that such a test could be devised by comparing the performance of QOM models on an experimentally-assembled network topology, with the performance of the same models re-estimated on networks whose topology has been appropriately randomized or “shuffled.” Importantly, such shuffles must carefully avoid possible confounds to their performance either because a shuffled network could *trivially* propagate more effects (e.g., by allowing more regulatory links) or because the shuffled topology would *effectively* allow more paths of propagation (at matched overall number of regulatory links) at higher orders. Figure 3A schematizes the simplest shuffle that controls for these effects by conserving the network topology while randomly re-assigning gene labels to its nodes; in effect, we synchronously randomize the rows and columns of the sparsity pattern for the regulatory matrix **B** before inferring its nonzero values during QOM model fitting (SI Appendix Sec. 6).

**Figure 3:**
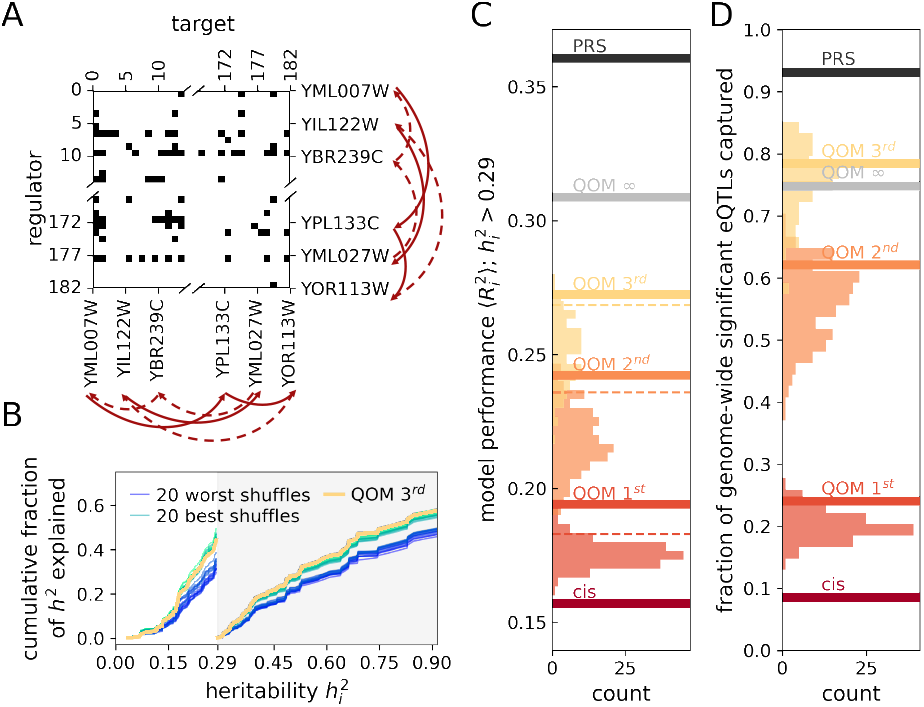
Experimentally-assembled GRN is more predictive than randomly shuffled networks. **(A)** Shuffling networks by re-naming the genes (i.e. simultaneously shuffling row and column labels) fully conserves network topology. **(B)** Cumulative sum of per gene fraction-of-variance explained, normalised by sum of heritability across genes, 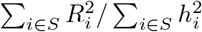, as a function of 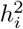, plotted separately for a set of genes *S* with 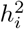below and above the median (0.29). Results shown for the 3^*rd*^ *order* QOM: thick yellow line corresponds to the experimentally-assembled GRN, thin green (blue) lines to the 20 best (worst) performing shuffled GRNs. The experimentally-assembled network outperforms most shuffled networks for 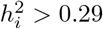 (grey shaded region). **(C)** Models based on experimentally-assembled GRN (solid lines) and shuffled GRNs (distributions). Shown is average per gene fraction-of-variance explained 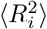, evaluated on genes with 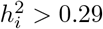. Number of shuffled networks for QOM 1^*st*^, 2^*nd*^ and 3^*rd*^ order QOMs is 111, 168 and 94, respectively. Dashed lines show 5% significance cuts in the matched shuffled distributions. **(D)** Fraction of GWS e-QTLs identified by Albert *et al* [1] captured by each model; plotting conventions same as in (C).

In line with our previous observations that QOMs show greatest potential for prediction improvement via effect propagation for genes with higher heritability (Fig 2A), we report no significant difference in fraction of heritable variance explained between the experimentally-assembled vs shuffled networks for genes with *h*^2^ *<* 0.29. In contrast, such a difference strongly and robustly emerges for genes *h*^2^ *>* 0.29 (plotted in Fig 3B for 3^*rd*^ *order* QOM). Figure 3C systematically tests the significance of network topology on such high-heritability genes, by comparing the experimentally-assembled network performance to the null distribution of performance obtained by fitting QOMs on randomly reshuffled networks. At each order, the experimentally-assembled-network QOM outperforms the shuffles at the *p <* 0.05 significance level (see Fig S8 for analogous analysis without the heritability cut). Lastly, we evaluated the fraction of genome-wide significant eQTLs captured by the shuffled networks, mimicking the last column of Fig 2D, showing that shuffles underperform the experimentally-assembled network also in this metric.

Randomizing a network while preserving its topology disrupts the correlation between *h*^2^ and gene in-degree (Fig 2E). To control for that, we constructed an alternative shuffle that conserves genes’ in-degree by construction. In this comparison, QOMs on the experimentally-assembled network outperform shuffled controls as well (Fig S8).

Next, the prior biological knowledge encoded in the details of regulatory network topology could be relevant for QOM performance only inasmuch as it reflects the functional grouping of genes. In this scenario, specific links between identified genes would not matter so long as the genes belonged to the same functional category. To test for that, we divided genes into “functional modules” based on their gene ontology biological function annotations [**GO2, GO1**], and re-assigned gene labels randomly within each module. Such GO-shuffled networks also under-performed the experimentally-assembled GRN (SI Appendix Sec. 6, Figs. S9A-C).

Indirect (*trans*-) effects arise by propagation of direct effects through the GRN. In the last control, we show that for all QOM orders, these network-selected *trans-* effects explain more variance compared to an equally large random set of markers with the same LD pattern (SI Appendix Sec. 6, Fig S9D).

In sum, the experimentally-assembled network QOMs outperform topology-conserving, in-degree-conserving, and functional-module-conserving shuffles, as well as a PRS on a random set markers. Performance gaps are higher for high *h*^2^ genes. This improvement cannot be explained solely by the larger number of free parameters (in-degree) for high *h*^2^ genes or a shared biological function of interconnected genes; rather, the regulatory network curated by the Yeastract+ consortium likely derives excess predictive power from its detailed pattern of connectivity that reflects actual molecular regulatory mechanisms. More generally, we put forward QOMs paired with shuffled controls as a comprehensive way of testing the quality of network reconstructions in a whole-genome, systematic manner. Importantly, such methodology could be deployed not only where full network reconstructions are available as in the case of yeast, but also for individual pathways or regulatory modules, at the scale currently more tractable for mammalian organisms, including humans.

### QOM predicts gene expression covariance

An individual mutation often has pleiotropic effects on multiple gene expression levels [41, 1, 18]. The omnigenic model rationalizes this observation and QOM turns it into a statistical model, which makes predictions beyond the marginal effects on individual genes. Specifically, the gene expression levels have a non-trivial correlation structure (Fig 4A): the distribution of pairwise correlation coefficients has long tails, with a large number of significantly correlated pairs whose correlation can extend up to |*c*_*ij*_ | *∼* 0.5 (Fig 4B), and which are predominantly arranged into salient rows or columns of the correlation matrix. Principal component analysis (Fig 4D, green line) reveals a much lower-dimensional underlying structure to these correlations, with 50% of the gene expression variance explained by the leading 8 principal components (PCs). We set out to test whether the observed correlation structure is captured by our models. Figure 4C evaluates the ability of the models to capture the observed gene expression correlations, by computing the *R*^2^ for the linear regression of the data vs model-predicted (off-diagonal) components of the correlation matrix. The 3^*rd*^ *order* model captures almost 40% of the variance in the correlation matrix elements and PRS about 50%. Higher-order QOMs are essential for this task: while the *cis* (0^*th*^ *order*) and the 1^*st*^ *order* models do a very poor job, fractionally large improvements in performance come from second-and third-order effect propagation – these improvements are much larger compared to the improvements in the prediction of individual expression levels (Fig 1E). To put the *R*^2^ numbers in perspective, we emphasize that we are comparing our (genetic) model predictions to the *total* gene expression correlation matrix, only part of which is genetic in origin and (possibly a substantial) part is environmental, as we further show in Fig 5.

**Figure 4:**
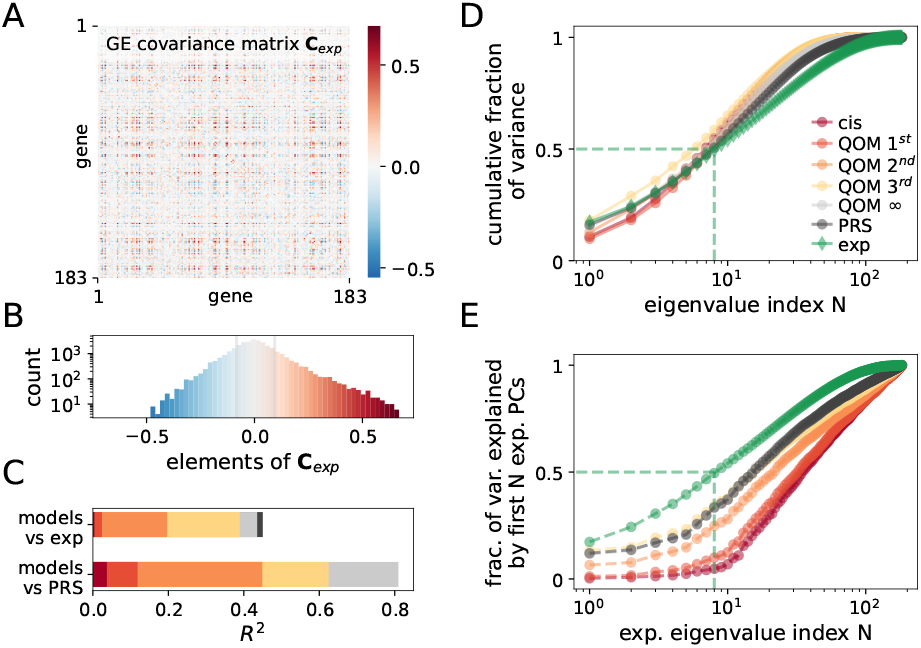
Models with higher-order effect propagation capture the covariance structure of gene expression. **(A)** Gene by gene covariance matrix (**C**_exp_) of experimentally measured expression levels **Y**_exp_ computed across individuals. Only off-diagonal elements shown for clarity. **(B)** Distribution of off-diagonal **C**_exp_ elements from (A). Shaded region contains non-significant correlations (*p* = 0.05 relative to shuffled data). **(C)** Fraction-of-variance (*R*^2^) for element-wise prediction by various models of data covariance (**C**_exp_, top bar) and of PRS predicted covariance (bottom bar). **(D)** Cumulative sum of ordered eigenvalues, for each model. 50% of variance in data is explained by the first 8 principal components (PC, dashed lines). **(E)** Overlap of PC subspaces between model predictions and data. Average squared fraction of the length of GE vectors predicted by various models lying in the data PC subspace spanned by its first *N* eigenvectors (green line is identical to that in D).

**Figure 5:**
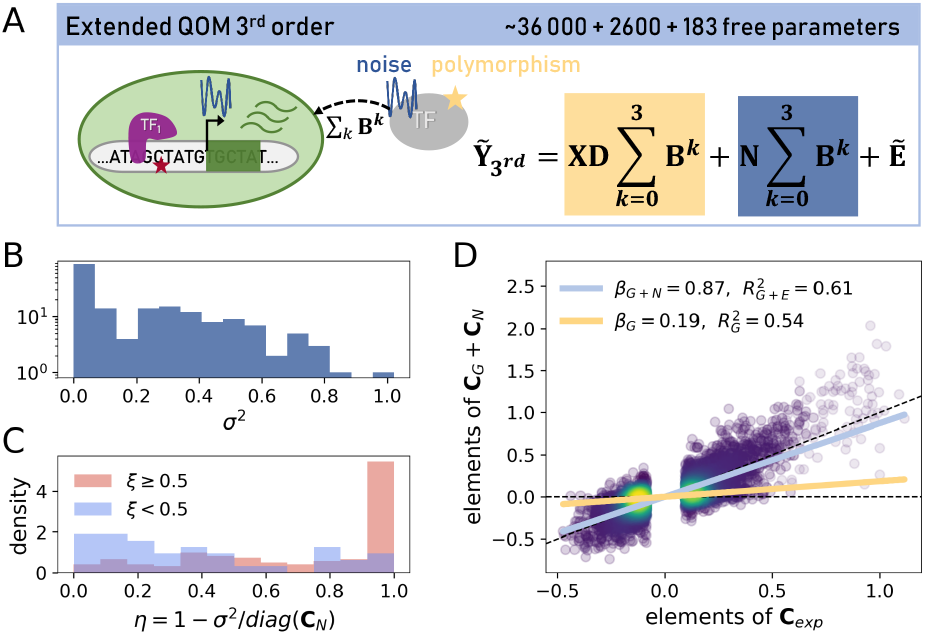
Transcriptional and non-transcriptional GE perturbations propagate through the same network. **(A)** Schematic of 3^*rd*^ *order extended* QOM for propagation of non-transcriptional effects **N** (dark blue) through the same network **B** that also propagates genetic transcriptional effects (yellow). **(B)** Histogram of inferred non-transcriptional effect variances,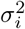, over *G* = 183 genes. **(C)** Distribution across genes of the fraction of propagated to total non-transcriptional variance *η* computed from inferred *σ*^2^ shown in (B). “Sensitive” genes dominated by *trans-* effects (*ξ* ≥0.5, red) tend to have higher *η* than genes dominated by *cis-* effects (*ξ <* 0.5, blue, SI Appendix Sec. 9). **(D)** Element-wise comparison of predicted genetic transcriptional (**C**_G_) plus non-transcriptional (**C**_N_) off-diagonal covariance to total GE covariance estimated from data (**C**_exp_), for gene pairs with significant non-zero covariance, as defined in Fig 4B. 3^*rd*^ *order extended* QOM predicts well also the magnitudes of GE covariance (blue linear regression, slope *β*_G+N_), compared to the plain 3^*rd*^ *order* QOM (yellow linear regression, slope *β*_*G*_); black dashed lines show zero (*β* = 0) and perfect (*β* = 1) GE covariance predictability, respectively.

Because the correlation matrix is dominated by a small number of eigendirections (Fig 4D), we asked to what extent various models reproduce the same structure. Figure 4E evaluates what fraction of variance in gene expression predicted by the QOMs and PRS lies in the subspace spanned by top-*N* PCs of the experimental GE, as a measure of the overlap of the respective PC subspaces. For the 3^*rd*^ *order* QOM, the QOM bound model, and the unconstrained PRS model – but not for lower-order QOMs – we see that the low-dimensional eigenvector subspace of the model strongly overlaps with the subspace identified by PCA directly on data. The variance in best performing models spanned by top 8 PCs constitutes around 38% of the total variance, or *≈* 75% of the experimental variance in the same subspace. This overlap is substantially higher than the element-wise comparison in Fig 4C would suggest, raising the possibility that a large share of environmentally or non-transcriptionally-propagated variation also resides in the same, genetically-identified subspace.

### Extended Quantitative Omnigenic model

To test this hypothesis, we extend the QOM to account for the propagation of perturbations to gene expression beyond those originating in identified *cis-*genetic effects: we call these extra sources of variation “non-transcriptional effects.”

These effects can either be (i) non-genetic, e.g., noise resulting from external environmental conditions or metabolic states which affect gene expression of other genes; or (ii) genetic yet non-transcriptional, e.g., due to genetic changes in the expression of non-transcriptional regulators of TFs, such as genes involved in metabolism, chromatin state, signalling, or modifying TF availability or activity. In fact, 68% of regulatory hotspots with effects on gene expression picked up by our PRS model do not overlap a TF [1]. Yet they affect gene expression of TFs, and consequently can propagate through the GRN.

The key idea is that a significant fraction of perturbations to gene expression that a population-level experiment can access, regardless of their origin, might propagate through the same regulatory network **B**, so that

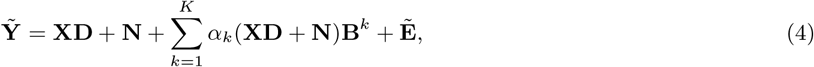

where **N** is the non-transcriptional effects matrix, **N** *∼ N* (0, **Σ**). **Σ** is a diagonal matrix with the non-transcriptional noise variance of each gene 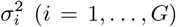 on the diagonal, to be inferred. If Eq (4) holds, residuals **E** in Eq (2) will not be independent of **B**: the residual *covariance* will have a structure given by the regulatory network matrix **B**, and 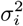 can be extracted from this residual covariance. This leads us to a 3^*rd*^ *order extended* quantitative omnigenic model, schematized in Fig 5A, which also propagates non-transcriptional effects at the increased complexity of *G* = 183 extra noise variance parameters, one per gene, which can be inferred by standard quadratic programming (SI Appendix Sec. 9).

Figure 2 suggests that the ratio of *cis-* and *trans-* GE variance varies across genes. In the extended QOM, we can ask how the fraction of genetic transcriptional variance in *trans* (*ξ*) compares to the fraction of non-transcriptional variance that is propagated via the network, *η*. We find that genes which have most genetic variance in *trans* (genes with *ξ ≥* 0.5), are also the genes that are dominated by propagated non-transcriptional effects, i.e., have high *η* (Fig 5C). This link between direct and propagated variance for transcriptional and non-transcriptional effects is suggested by the omnigenic model under certain assumptions (SI Appendix Sec. 9), motivating further analysis.

We observe that while the data-estimated covariance elements correlate well with those predicted by 3^*rd*^ *order* QOM (Fig 4C), the absolute magnitude of covariance in the data is much larger than predicted (Fig 5D, yellow line). Because covariance elements are pure predictions of QOMs with no free parameters, these predictions indeed can (and do) correlate well with data-derived covariance estimates even as they severely underestimate their absolute magnitude. This particular structure of deviations from data is consistent with our hypothesis that non-transcriptional sources of variability likely propagate through the same regulatory network, and the 3^*rd*^ *order extended* QOM successfully captures such propagation (Fig 5D). Specifically, while the linear correlation coefficient between GE covariance element predictions and data estimates improves only marginally from the 3^*rd*^ *order* QOM to its *extended* version (increasing from linear correlation of 0.73 to 0.78, respectively), the best-fit linear slope improves drastically from 0.19 to 0.87, closing most of the covariance magnitude gap. In short, QOMs have the potential to explain the propagation of direct genetic *as well as* non-transcriptional genetic and non-genetic (noise) effects, to successfully account for the total – genetic plus environmental – structure of gene expression covariance. Methodologically, our findings further suggest that improved statistical methods for GWAS could quite generally make use of the fact that noise **E** in Eq (1) appears structured, as we show above.

## Discussion

We introduced the Quantitative Omnigenic Model (QOM), a statistical counterpart and a formalization of the omnigenic hypothesis [6]. QOM is a variant of GWAS that incorporates known prior knowledge about intracellular networks to propagate effects of mutations that affect observed traits only over experimentally confirmed network links. By doing that, prior knowledge imposes very strong structural and parameter constraints on QOM, compared to an unconstrained GWAS-type regression such as a polygenic risk score (PRS) model. It should therefore be clear that the primary aim of QOM is not to outperform GWAS or PRS-type models in raw trait predictability, but rather complement it by dissecting how intracellular networks mediate *trans-* and *cis-* effects, and, in the process, generate compact causal and mechanistic hypotheses about pathways of effect propagation and the overall architecture of gene expression traits.

To demonstrate the breadth of interpretable results accessible within the QOM framework, we applied it to a yeast cross dataset with gene expression phenotype measurements [1]; mutational effects were propagated through the *S. cerevisiae* transcriptional regulatory network as curated by the Yeastract+ consortium [25]. We report multiple findings relevant to genomics and, specifically, to the architecture of gene expression traits. **First**, QOMs reach performance competitive with unconstrained PRS models despite orders-of-magnitude less free parameters, giving credence to approaches that incorporate biological knowledge into statistical genomics models *a priori* rather than using it solely *post hoc*. **Second**, QOMs confirm that 50% of heritable variance in gene expression in the analyzed dataset can be explained by (up to) three steps of propagation of mutational *cis-* effects through the transcriptional GRN, in line with the omni-genic hypothesis. **Third**, a comparison of *cis* vs 3^*rd*^ *order* QOM suggests that at least *∼* 30% of heritable GE variance is in *trans* in a way that QOMs can trace via explicit pathways of propagation; a further *∼* 15% may be reachable via more steps according to the QOM bound which itself is within five percent of PRS. Fraction of heritable variance due to transcriptional effects on GE in *trans* is thus 1.5 *−* 2*×* that in *cis*. **Fourth**, a comparison of PRS vs 3^*rd*^ *order* QOM identifies a small number of yeast TF genes (7), as well as a substantial fraction of previously reported eQTLs (51%) that are not explained by or do not predict GE via transcriptional effect propagation, respectively. Together, these interactions are responsible for the gap between the QOM and PRS that accounts for 6% of gene expression variance, likely attributable to non-transcriptional *trans* effects. **Fifth**, the analysis of individual pathways of effect propagation highlights the preponderance of *omnigenes* and suggests that polygenicity for GE is even more widespread than traditional GWAS would suggest. **Sixth**, QOM reveals correlations between heritability and predictability of GE and the in-degree of the corresponding genes, raising interesting questions about the traits’ mutational robustness, tunability, and evolvability. Exploration of other potential links between GE predictability and local network topology is a promising topic for future research.

Two further findings are of high relevance to systems biology. **First**, we demonstrate that QOMs can be used as a statistical test for the quality of regulatory network reconstructions. While such reconstructions could be tested link-by-link with costly mutational, pharmacological, or environmental interventions, we are unaware of any systematic quantitative procedure that would simultaneously assess the entire network topology. Instead of targeted, costly interventions, here we used polymorphisms generated in a genomic experiment as a massive, distributed, and weak probe of gene expression levels. If the reconstructed network topology reflects the underlying regulation mechanisms, it should be more predictive in a statistical sense compared to randomized networks when tested on polymorphism data. We confirm that this is indeed the case for the yeast GRN and suggest that such testing methodology is applicable to other networks, pathways, and regulatory modules. In general, this network test serves as a guide of QOM applicability: whenever a given pathway or network is more predictive than appropriate randomised controls, the QOM is picking up real signal. **Second**, while genomics focuses on propagation of genetic effects, often assuming a trivial structure for “environmental noise,” systems biology mostly focuses on the propagation of external effects (nutrients, chemical signals, metabolic state, environmental variables) while holding the genotype fixed. Yet it is reasonable to hypothesize that *both* types of effects propagate along the same intracellular networks. This was the motivation for the *extended* QOM that supports our hypothesis, by successfully capturing the structure of the *total* – i.e., direct genetic, non-transcriptional genetic and environmental – gene expression covariance matrix.

We set out to explore the system-wide implications of the omnigenic hypothesis and, therefore, decided to apply the QOM framework to a system for which the entire regulatory network has been reconstructed, the inferences are tractable, and benchmark comparisons exist. This restricted our choices and, correspondingly, the generality of our findings. Our goal was to assess whether assembled regulatory networks carry predictive power and can, in principle, be utilised in interpretable predictive models. Therefore, we used a dataset with maximal predictive power: a yeast cross with MAF*∼* 0.5 and block-like LD pattern (Fig S10). This scenario is ideal because direct effects can be detected reliably and are not linked to *trans-*eQTLs. Propagation of these direct effects via transcriptional pathways can further suggest causal *trans-*eQTLs. Further, our data come from a fixed environment, i.e., we do not observe variation in expression due to changing environments. Moreover, there are more eQTLs segregating in wild yeast populations [24] than Albert *et al* report [1]. As a consequence, the reported fraction of GE variance explainable by transcriptional regulation is likely specific to our data, and should in general depend on population and environmental conditions.

To explore the applicability of QOM to populations more representative of the standing variation, we deployed QOM in a supplementary analysis on yeast “pan-population” dataset composed of natural *S*.*cerevisiae* isolates from various ecological and geographical niches [**Caudal2024**] (Fig S10). We find that QOM successfully predicts gene expression and suggests causal transcriptional regulatory interactions also on this data, despite the notably lower statistical power compared to the cross used in our main analysis (Fig S11, SI Appendix Sec. 11).

Despite its limitations, this work showcases the potential of employing traditional statistical genomics models side-by-side with structured models that incorporate prior knowledge about regulatory networks in order to flexibly trade off interpretability with prediction performance. Paying careful attention to statistical tests of QOM validity using randomized networks as we propose, QOM could be attempted on further natural populations. Another possibility would be to restrict its application to individual pathways or network modules in other organisms – especially if these pathways relate to selected phenotypes for which more reliable prior information may be available, e.g., on cellular processes or metabolism. To what extent QOM can successfully be utilised for interpretable predictions on statistically more challenging datasets beyond the two study yeast populations we focused on here remains the subject of future work.

## Materials and Methods

The main analysis was conducted on a population of 1012 haploid segregants from BYxRM yeast cross by Albert et al genotyped on 42 052 and gene expression measurements for 5724 genes. Training was done on 800 individuals and the remaining 212 were used for model evaluation. The GRN topology for the main analysis was defined as the 183 *×* 183 subnetwork of TF genes of the Yeastact+ DNA binding evidence network assembled from chromatin immunoprecipitation (ChIP), ChIP-on-chip, ChIP-seq and electrophoretic mobility shift assay studies in *S*.*cerevisiae* [25]. The detailed materials and methods are described in the SI Appendix. The code used for fitting and evaluating the QOM as well as shuffling regulatory networks is available at https://github.com/nataliaruzickova/Quantitative-Omnigenic-Model.git.

## Acknowledgements

NR acknowledges the support of the Austrian Academy of Sciences through the DOC fellowship 26917. MH and GT were supported in part by the Human Frontiers Science Program Grant RGP0034/2018. We thank Nicholas H Barton, Fyodor Kondrashov, and Matthew R Robinson for fruitful discussions.

