## Supplementary information and figures for "Quantitative omnigenic model discovers interpretable genome-wide associations"

### Supplementary Information for: Quantitative omnigenic model discovers interpretable genome-wide associations

The manuscript was compiled on September 28, 2024

#### This PDF file inculdes:

SI Appendix
Supplementary figures S1-S11

*Institute of Science and Technology Austria, AT-3400 Klosterneuburg, Austria

### SI Appendix

#### 1 Models

**Training and error estimates.** We train all models on 800 randomly chosen individuals (train set) and use the remaining 212 for hyperparameter optimization and model performance evaluation (evaluation set). We use Lasso (L1-norm) regularisation in all models, the strength of which ($\lambda$) is a hyperparameter for each model determined by sub-sampling in the following way: in each of 50 folds, we randomly choose 100 segregants from the evaluation set and compute the average performance across folds for each value of $\lambda$. Then we choose $\lambda$ which maximises this performance, and evaluate the performance on the remaining 112 individuals in each fold. The average (standard deviation) across folds of this test performances is the reported performance (errorbar) in Fig 1E. By doing this evaluating on 50 pairs of sub-samples of the evaluation set, we i) reduce the bias of randomly choosing only one set of 100 segregants for validation and the remaining 112 for testing and ii) estimate the error resulting from using a random subset of the population for evaluating our models. These errors are rather small, as Fig 1E shows.

**Performance measures.** We report multiple measures of performance: (1) total fraction-of-variance explained across all TFs, $R^2$; (2) average per gene (gene by gene) fraction-of-variance explained, $\langle R_i^2 \rangle$; and (3) average per gene fraction-of-heritable-variance explained, $\langle R_i^2/h_i^2 \rangle$. In all cases $R^2$ is the squared Pearson correlation between experimentally measured and model predicted expressions for a subset of individuals, $R^2 = r^2(y_{model}, y_{exp})$. (1) is used for training of the models.

**Structure of the models and biological evidence.** Overall, we study 6 models: QOM $0^{th}$ *order* (*cis*, (M2)), QOM $1^{st} - 3^{rd}$ *order* (M1), QOM bound model (QOM $\infty$, (M7)) and unconstrained "Polygenic risk score-type" model (referred to as PRS, (M9)). QOM model of order $K$ is of form, as defined in (2) with $\alpha_k = 1$:

$$\mathbf{Y} = \mathbf{XD} + \mathbf{XD}\sum_{k=1}^{K}\mathbf{B}^k + \mathbf{E} \tag{1}$$

where $\mathbf{X}$ are genotypes, $\mathbf{Y}$ are gene expression levels (see SI Appendix Sec. 2 for precise definition), $\mathbf{D}$ is the direct effects matrix (*cis*-eQTLs) and $\mathbf{B}$ is the transcriptional regulatory network. QOM models are constrained by two types of biological evidence:

1. $d_{ij}$ is 0 whenever polymorphism $i$ is not within a confidence interval around a *cis*-eQTL to gene $j$, as reported by Ref [1] (Fig SM1B). The nonzero entries of the $P \times G$ matrix $\mathbf{D}$ (direct effects on gene expression) are determined by fitting QOM $0^{th}$ *order*.

2. entry of the regulatory network matrix $b_{ij}$ can be nonzero when $i \neq j$ (no self loops) and when there is experimental evidence of regulatory interaction between regulator $i$ and target gene $j$ in database Yeastract+ [4]. Yeastract+ lists two types of underlying experimental evidence for a regulatory in *Saccharomyces cerevisiae*, which the authors define as

   (a) *DNA binding evidence*: "chromatin immunoprecipitation (ChIP), ChIP-on-chip, ChIP-seq and electrophoretic mobility shift assay studies;"

   (b) *Expression evidence*: "data obtained from the comparative analysis of gene expression changes in response to the deletion, mutation or over-expression of a given TF (based on reverse transcriptase-polymerase chain reaction, microarray analysis, RNA-seq or expression proteomics)."

   The structure of both networks is shown in Fig. fig. S3. We choose the GRN defined by *DNA binding evidence* (Fig SM1C) for main analysis since *Expression evidence* may by design include links which are not a causal direct regulatory interaction, but a propagated interaction through an intermediate gene.

   In the 183x183 *DNA binding evidence* GRN, all TFs apart from `YPR199C` have a nonzero in-degree, which means they are regulated by at least one of the other TFs. 35 TFs do not have targets among the 183 TFs (zero out-degree). We replicate the key analysis for the *Expression evidence* in Fig. S2 and SI Appendix Sec. 7. All remaining analysis is done using *DNA binding evidence* based GRN, unless stated otherwise.

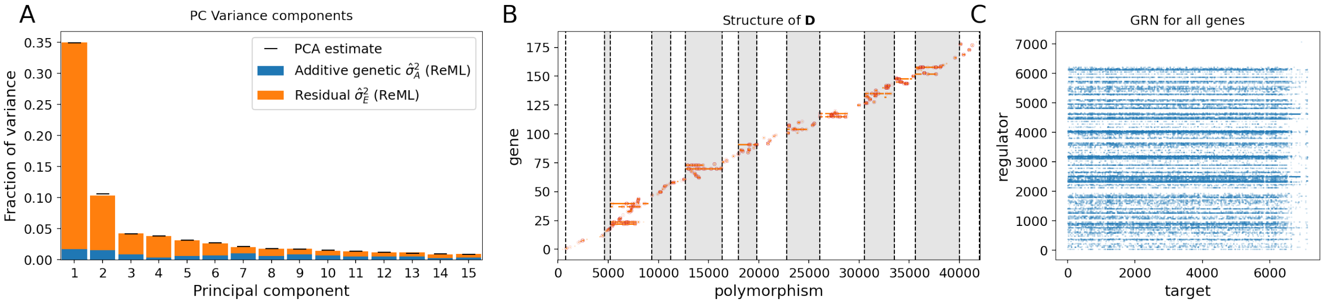


Figure SM1: **Preliminaries. (A)** Variance explained by successive principal components of gene expression (line segments; see legend). Colored bars show variance components of GE PCs estimated using maximum likelihood (ReML; PCA done on TF genes, corrected for batch and OD). The heritability of PC1 is very low - less than 5%. Therefore it is likely that PC1 corresponds to other technical or metabolic variation (possibly variable growth rate / nutritional uptake, as shown for example by [2]). **(B)** Structure of the direct effects matrix **D** based on *cis*-eQTLs and their confidence intervals reported by Ref [1]. Out of all 42 052 polymorphic sites, only those which are within a confidence interval of a *cis*-eQTL to one of the 183 TF genes (19 720 sites, 47%) can contribute in QOMs (orange dots). L1 regularisation in the *cis* model leaves only 377 of them nonzero (red dots), implying that the variance in expression of all genes of interest is explained solely by these 377 sites in the QOMs. **(C)** Transcriptional GRN of all genes from Yeastract [4] based on DNA binding evidence. Rows with many non-zero elements (blue) correspond to TFs. This structure is characteristic for transcriptional networks and it strongly influences the properties of QOMs.

**QOM $0^{th}$ *order* (*cis*) model.** We fit QOM $0^{th}$ *order* separately for each gene $i$ by linear regression with L1 penalty (Lasso, python scikitlearn implementation). This is essentially multivariate multiple regression:

$$(y_{cis})_i = \mathbf{X}^{(\mathbf{i})} d_i + e_i, \tag{2}$$

where $d_i$ is the vector of *a priori* nonzero entries of the $i^{th}$ column of $\mathbf{D}$, $\mathbf{X}^{(\mathbf{i})}$ is the matrix $\mathbf{X}$ with genotypes of only those polymorphisms which are in *cis* to gene $i$, and $e_i$ are the residuals. Matrix $\mathbf{D}$ obtained from this regression is used further in QOM $1^{st} - 3^{rd}$.

**QOM $1^{st}$ *order* model.** QOM $1^{st}$ *order* is fitted analogously to QOM $0^{th}$ *order*, by Lasso linear regression on *a priori* nonzero entries of the GRN matrix $\mathbf{B}$ (i.e., $b_{ij} = 0$ if there is no regulatory interaction detected between regulator $i$ and target $j$ according to Ref [4]):

$$(y_{1^{st}})_i - \mathbf{X} d_i = (\mathbf{XD})^{(\mathbf{i})} b_i + e_i, \tag{3}$$

where $b_i$ and $d_i$ are the $i^{th}$ columns of matrices $\mathbf{B}$ and $\mathbf{D}$, respectively. $b_i$ is fitted here, $d_i$ is fixed to the value obtained from QOM $0^{th}$ *order* model fit and $(\mathbf{XD})^{(\mathbf{i})}$ is a matrix with columns corresponding only to genes which regulate gene $i$.

**QOM $2^{nd}$ and QOM $3^{rd}$ *order* models.** QOM $2^{nd}$ and QOM $3^{rd}$ *order* are non-linear models fitted using numerical L-BFGS-B method [3, 5] (python implementation) derived from BFGS (Broyden–Fletcher–Goldfarb–Shanno) algorithm, which is a numerical quasi-Newton method that estimates the inverse Hessian. L-BFGS-B uses limited memory and bounds on the values of the unknown vector.

We use L-BFGS-B to fit the *a priori* nonzero entries of the $\mathbf{B}$ matrix for $K = 2$ and $K = 3$:

$$\mathbf{Y}_k - \mathbf{XD} = \mathbf{XD} \sum_{k=1}^{K} \mathbf{B}^k + \mathbf{E}. \tag{4}$$

For convenience, we define the propagated effects network which includes all direct and indirect regulator-target interactions (all *trans*-eQTLs) up to order $K$ possible under regulatory network $\mathbf{B}$ :

$$\tilde{\mathbf{B}}_K = \sum_{k=1}^{K} \mathbf{B}^k. \tag{5}$$

Since we are interested in selecting only those regulatory interactions which are contributing significantly out of the set of possible ones given by Ref [4], we initialize all regulatory links at 0, and let the model pick those which are nonzero while using (a smooth relaxation of) L1 regularization.

Because the smooth L1 / Lasso did not push the individual entries strictly to 0 but left some at tiny nonzero values, we chose a cutoff of $10^{-3}$ for the magnitude of entries of $\mathbf{B}$ for QOM $1^{st} - 3^{rd}$ *order*. The cutoff for total genetic effect magnitude of QOM model of order $K$, i.e.,

$$\mathbf{G}_K = \mathbf{D} + \mathbf{D}\tilde{\mathbf{B}}_K \tag{6}$$

for the QOM bound model, as well as PRS model, was set to $10^{-11}$. Both thresholds were chosen based on a clear gap in the distribution of effect sizes of regulatory links and in the distribution of genetic effects, which was consistent across models, as shown in Fig SM2. After fitting the models, entries of matrices $\mathbf{D}$, $\mathbf{B}_K$ and $\mathbf{G}_K$, below the corresponding threshold were set to zero, and further analysis was done with such thresholded matrices.

**QOM bound / QOM $\infty$ model.** We define an upper bound to the QOM models in the following way. We fit a PRS-type model

$$\mathbf{Y} = \mathbf{X}\mathbf{G}_{\text{QOM}} + \mathbf{E}, \tag{7}$$

where the nonzero *sparsity structure* of $\mathbf{G}_{\text{QOM}}$ is given by

$$\bar{\mathbf{G}}_{\text{QOM}} = \bar{\mathbf{D}}\left(\mathbb{1} - \bar{\mathbf{B}}\right)^{-1} \tag{8}$$

where $\bar{\mathbf{A}}$ is the prior sparsity structure of matrix $\mathbf{A}$, i.e., a matrix of 0/1 elements, where 1 stands for the entries that are fitted by the model, and 0 stands for entries that remain fixed to 0. $\bar{\mathbf{D}}$ is shown in Fig. SM1B (orange dots) and $\left(\mathbb{1} - \bar{\mathbf{B}}\right)^{-1}$ in Fig. S3F. We then fit the QOM bound model gene-by-gene using L1-penalised linear regression, analogously to matrix $\mathbf{D}$ in QOM $0^{th}$ *order* described in Section 1.

We interpret Eq. (7) through a Taylor expansion $(1 - \mathbf{B})^{-1} = \mathbb{1} + \sum_{k=1}^{\infty} \mathbf{B}^k = \mathbb{1} + \tilde{\mathbf{B}}_\infty$. Qualitatively, the structure of $\mathbf{G}_{\text{QOM}}$ (Fig S3) tells us which sites can potentially be *trans*-eQTLs to which genes after $\infty$ number of propagation steps. Note that $\bar{\mathbf{G}}_{\text{QOM}}$ is not a full matrix: around 59% of its entries are zero, because of the nontrivial structure of the $\mathbf{B}$ matrix (rows corresponding to master regulator genes are nearly full).

We note that QOM bound model is not equal to QOM of order $K = \infty$ in the mathematical sense as defined by (M1) (one marker effect = linear combination of direct effects and regulatory links). Instead, the QOM bound model is a "PRS-type" linear model (one marker effect = one free parameter), where the *sparsity* of $\mathbf{G}_{\text{QOM}}$ is given by *structure* of $\mathbf{D}$ and $\mathbf{B}$ and the nonzero entries of $\mathbf{G}_{\text{QOM}}$ are fitted *independently* (i.e., not computed from $\mathbf{B}$ and $\mathbf{D}$ ). QOM bound model differs from PRS model only in its *a priori* defined sparsity structure. In the text, we use the notation "QOM bound" and "QOM $\infty$" interchangeably.

**Polygenic Risk Score (PRS)-type model.** "PRS" in our paper is an unstructured and unconstrained L1-regularised linear regression model:

$$\mathbf{Y} = \mathbf{X}\mathbf{G} + \mathbf{E}, \tag{9}$$

where any element of $\mathbf{G}$ can be nonzero, and the sparsity pattern is determined solely by regularisation.

### 2 Data

To train our models we use data provided by Albert *et al* [1] on 1 012 haploid segregants from a yeast cross between laboratory (BY) and wild (RM) strain. All individuals were genotyped at 42 052 polymorphic sites, which covers most of the genetic differences between the two strains. Expression levels of 5 724 genes were measured for all segregants divided into 13 batches. The expression levels (measured in units of $\log_2$ TPM - Transcripts per Million) were further corrected for technical covariates (OD and batch), standardised and corrected for the first principal component which captures non-genetic – likely technical or metabolic – variation (Fig SM1A), and centered.

The genotypes were normalised by allele frequency $X_{ij} = \frac{x_{ij} - p_j}{\sqrt{p_j(1-p_j)}}$ where $x_{ij}$ is either 0 (reference allele) or 1 (alternative allele) at locus $j$ of individual $i$ and $p_j$ is the frequency of alternative allele. We focus our analysis on transcription factors, which we define as genes that, based on *DNA binding evidence* (see Section 1), according to the Yeastract+ database [3], regulate other genes. There are 184 such genes, for three of which (`YJL089W, YLR013W, YSC0015`) there are no expression measurements by [1]. To keep the GRN as intact as possible, we replaced the expressions of YJL089W and YLR013W with a gene with most similar expression pattern, according to the SPELL database [2], which are YML042W and YOL015W, respectively. Neither of them is a TF, so all GE vectors in the $\mathbf{Y}$ matrix are unique. SPELL provides no information on YSC0015, therefore we excluded it from the analysis, leaving us with 183 TF genes.

**LD correction.** We defined discrete LD blocks in the genome as follows. First, for each chromosome, we compute genetic correlation $\rho$ from the genotypes matrix of all individuals and all markers. Then we estimate the number of blocks in each chromosome such that the mean LD within blocks is around 0.95: the number of blocks and the mean LD for each chromosome are indicated in Table1. Then, we use K-means clustering to cluster markers into the defined number of clusters on each chromosome, and each cluster is an LD block.

| chromosome | I | II | III | IV | V | VI | VII | VIII | IX | X | XI | XII | XIII | XIV | XV | XVI |
|---|---|---|---|---|---|---|---|---|---|---|---|---|---|---|---|---|
| # LD blocks | 13 | 25 | 15 | 35 | 23 | 13 | 32 | 23 | 20 | 30 | 25 | 32 | 30 | 29 | 35 | 35 |
| mean LD | 0.97 | 0.94 | 0.95 | 0.94 | 0.96 | 0.95 | 0.95 | 0.97 | 0.96 | 0.96 | 0.94 | 0.94 | 0.95 | 0.97 | 0.96 | 0.96 |

Table 1: **Number of LD blocks and mean LD at each chromosome.**

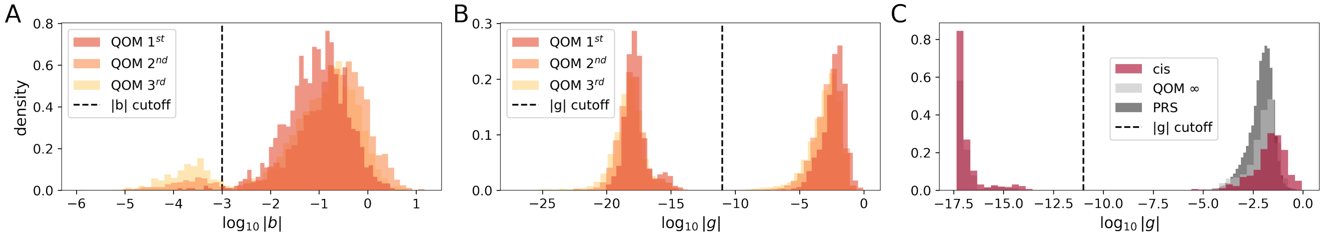


Figure SM2: **Regularization-induced gap in the distribution of inferred effect sizes.** Distribution of absolute values of effect sizes before applying any cutoffs and the corresponding cutoff thresholds for **(A)** regulatory strengths (dashed vertical cutoff line at $10^{-3}$) and **(B, C)** genetic effects (dashed vertical cutoff lines at $10^{-11}$).

### 3 ML upper bound variance estimates

To upper bound the variance explained by regression-like QOM and PRS models, we used an independent maximum-likelihood (ML) variance component estimation method. In the main text we refer to these estimations as "ML bounds". The goal is to estimate the variance in gene expression explained by a set of markers identified as *cis*- and *trans*-eQTLs by the QOM (based on an experimentally informed GRN), and compare the amount of variance explained by different orders of propagation.

**Standard estimation of two variance components.** The standard way to decompose variance in a trait is using a linear model [2],

$$\vec{y} = \mu\vec{1} + \vec{a} + \vec{e}, \tag{10}$$

where $\vec{y} = (y_1, \ldots, y_N)^T$ is a vector of trait values for $N$ individuals, in our case log transformed and standardised expression levels corrected for technical covariates and PC1, as described in Section 2. On the right hand side, $\mu$ is the mean and $\vec{1}$ is a vector of ones, $\vec{a}$ are the additive genetic values and $\vec{e}$ is the vector of residuals – biological noise, non-additive genetic effects, environmental and unaccounted-for technical effects and measurement errors.

This is a statistical mixed model where the measured values of $\vec{y}$ and genotypes $X \in \{0,1\}^{N\times P}$ (for $P$ haploid marker loci) are available. The residuals are assumed to be uncorrelated with unknown variance $\mathrm{Var}[\vec{e}] = \sigma_E^2 \mathbb{1}$. The additive genetic component $\vec{a} \sim \mathcal{N}(0, \sigma_A^2 \mathbf{A})$ has a covariance matrix given by the unknown additive genetic variance $\sigma_A^2$ and the known (additive) relatedness matrix between measured individuals,

$$\mathbf{A} = \frac{N\mathbf{X}'\mathbf{X}'^T}{\mathrm{Tr}\left(\mathbf{X}'\mathbf{X}'^T\right)}, \tag{11}$$

where $\mathbf{X}'$ is the centered matrix of genotypes; in haploids $X'_{ij} = X'_{ij} - p_j$ where $p_j$ is the frequency of allele 1 at locus $j$. This formula for the relatedness matrix $\mathbf{A}$ can be derived by assuming that the additive effects at all loci are random with a constant variance.

Both $\vec{a}$ and $\vec{e}$ are assumed to have a multivariate normal distribution, leading to a distribution of $\vec{y}$,

$$\vec{y} \sim \mathcal{N}(\mu\vec{1} + \sigma_A^2 \mathbf{A} + \sigma_E^2 \mathbb{1}), \tag{12}$$

which serves as a likelihood function. The goal is to estimate the two variance components $\sigma_A^2$ and $\sigma_E^2$ while accounting for the unknown mean $\mu$.

A standard method is restricted maximum likelihood (ReML)[2], which is a variant of maximum likelihood inference applied within a subspace of $\vec{y}$ orthogonal to subspace spanned by the vector $\vec{1}$. As the maximisation method for ReML we use EMMA [1]. EMMA speeds up likelihood evaluation by cleverly storing the decomposition of suitable matrices.

This ReML algorithm is used to estimate the heritabilities of different principal components in Fig SM1A.

**Variance explained by a subset of polymorphisms.** Since we want to partition variance between two sets of polymorphisms (model-selected vs the rest), we split the additive component into two parts:

$$\vec{z} = \mu\vec{1} + \vec{a}_1 + \vec{a}_2 + \vec{e}, \tag{13}$$

where the additive components 1 and 2, $\vec{a}_1$ and $\vec{a}_2$, respectively, are associated with two different relatedness matrices, $\mathbf{A}_1$ and $\mathbf{A}_2$, each computed from the corresponding subset of markers, and two variance components, $\sigma_1^2$ and $\sigma_2^2$.

We obtain the ReML estimate of $\sigma_1^2$, $\sigma_2^2$ and $\sigma_E^2$ by a modified EMMA algorithm [1] (self-implemented in python). For a fixed fraction of additive variance in the first set of markers, $\rho = \frac{\sigma_1^2}{\sigma_1^2 + \sigma_2^2}$, maximisation can be done by EMMA. The maximum across different values of $\rho$ is found by a one-dimensional grid search followed by a binary search. The calculation returns the ReML estimates $\hat{\sigma}_1^2$, $\hat{\sigma}_2^2$ and $\hat{\sigma}_E^2$.

The calculation is done separately for each trait (expression of each gene). As input, we need to specify the criterion for markers in group 1 vs in group 2. For each gene and a given model (QOM or PRS), the markers in group 1 are those that the model selected to have a non-zero effect on the given gene's expression, plus their LD blocks (for LD correction see Section 2).

We note that in this upper bound estimation, the structure of the GRN is reflected solely in the *set of markers* which can potentially have a non-zero effect on a gene's expression, similarly to QOM bound model explained above. Each marker within this set is treated *independently* of the others, in contrast to QOM 1-3 ((M3), (M4)) which simulate propagation of effects through the given network. Due to this distinction, the ML bound is not a tight upper bound.

### 4 Consistency measures

**Consistency between QOMs.** To assess coherence between models, we measure whether pathways of effect propagation have been identified consistently by different QOMs. Specifically, if QOM of order $K = 2$ identifies regulators (*trans*-eQTLs) of a given gene and their associated regulatory weights, we the asses how well these upstream effects align with those identified by an independent QOM of order $K' = 3$ following, for instance, $k = 1$ step of propagation. Consistency between models of order $K$ and $K'$ for a given gene is achieved when these two models consistently pinpoint the same regulatory pathways upstream of that gene. In simpler terms, it means that the set of regulators and their relative contributions match. Consequently, the *trans*-eQTL effects associated with these regulators cluster along a line. Therefore, as a measure of consistency, we use Pearson correlation between the two pathways. Examples in Fig SM3A have high consistency.

Mathematically, we define consistency between QOMs of order $K$ and $K'$ at distance $k$, $k \leq min(K, K')$ for regulatory pathways of gene $i$ as

$$O_{(K,K')}^{(k)}(i) = r\left([\mathbf{D}\mathbf{B}_K^k]_{:,i}, [\mathbf{D}\mathbf{B}_{K'}^k]_{:,i}\right). \tag{14}$$

Here, $\mathbf{D}$ is the direct effects matrix inferred by the *cis* model (M2), $\mathbf{B}_K$ is the GRN matrix inferred by QOM of order $K$ (M1), $r$ is Pearson correlation, which we compute on propagated genetic effects from regulators of gene $i$ (the $i^{th}$ column of $(\mathbf{D}\mathbf{B}_K^k)$ for QOMs of order $K$ and $K'$). We compute consistency independently for each gene $i$ and for each set of parameters $K, K'$ and $k$.

We measure consistency only for the pathways which are in principle identifiable / reachable by the lower-order model. We only consider whether the regulatory interaction strengths inferred by two QOMs cluster along a line (are proportional), not whether the magnitudes match, since the number of propagation terms in QOMs varies, and the magnitude of terms on inferred **B** varies as well. Illustration for one example gene is shown in Fig SM3A, and distribution of consistency scores across individual genes is shown in Fig SM3B. Finally, to report a *single* consistency score per gene (Fig 2D and Fig S5), we average the consistency between each pair of models:

$$\text{consistency}(i) = \frac{1}{4}\left[O^{(1)}_{(1,2)}(i) + O^{(1)}_{(2,3)}(i) + O^{(1)}_{(1,3)}(i) + O^{(2)}_{(2,3)}(i)\right]. \tag{15}$$

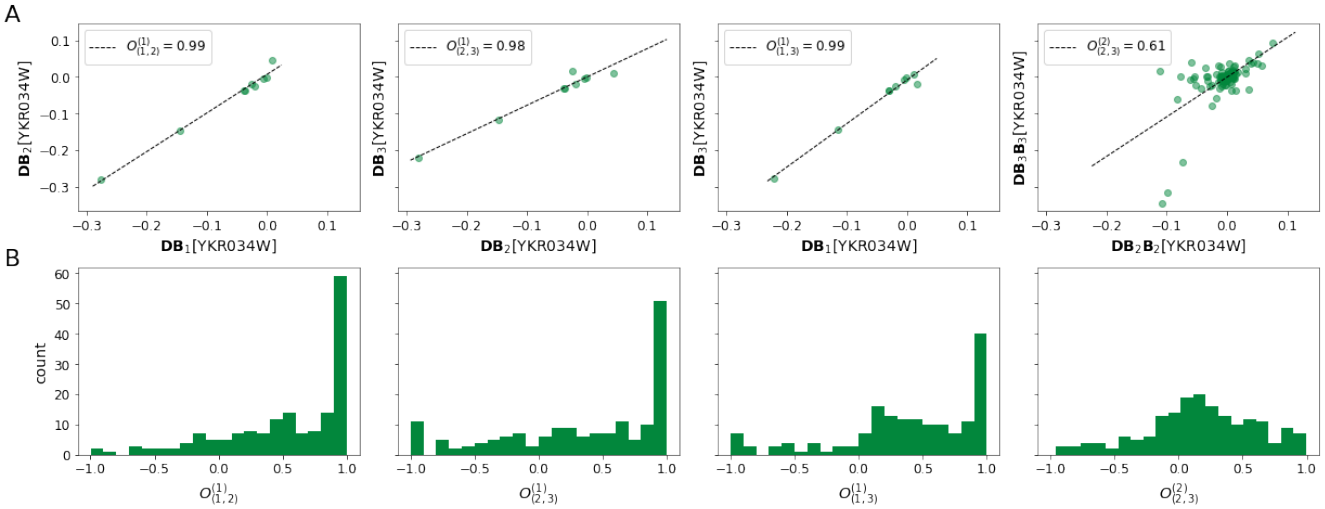


Figure SM3: **Consistency at which QOMs identify regulatory pathways.** **(A)** Definition of consistency between individual QOM orders according to Eq. (M14), illustrated for the example gene `YKR034W` for all combinations of $k \in \{1, 2\}$ and $K, K' \in \{1, 2, 3\}$ in for gene `YKR034W`. Each dot is one propagated genetic effect (*trans*-eQTLs of `YKR034W`). On the axes are sizes of these effects, as inferred by two different QOMs. First three plots compare first order of propagation ($k = 1$ in (M14)) for the three QOM pairs ($K, K' \in \{1, 2, 3\}$), the fourth plot shows second order ($k = 2$) as inferred by QOM2 vs QOM3 ($K = 2, K' = 3$).The dashed line is a best linear fit (with $r = O^k_{K,K'}$ indicated in the legend) serving as a guide to the eye. **(B)** Distribution of consistency scores defined by (M15) across all 183 genes.

**Consistency with previously reported genome-wide significant (GWS) e-QTLs.** As a further consistency measure we look at the overlap between eQTLs identified by our models (both *cis* and *trans*) and GWS eQTLs reported by Ref [1] (on the same dataset, but obtained by an unrelated method). Specifically, we look at the *fraction of GWS eQTLs reported by Ref [1] picked up by our model*: the number of GWS eQTLs reported by Ref [1] which are among QOM/ PRS sites with nonzero effect, divided by the total number of these GWS eQTLs. Results for all genes are shown in Fig S5B,D (last column). Fig SM4 shows the match between the signs of effects of GWS eQTLs detected by Ref [1] and those detected by our models, averaged through LD blocks. The entries on the diagonals denote the fraction of model-detected effects whose sign matches the GWS eQTL sign. We note that our normalization of gene expressions differed from the original study, therefore even for PRS we do not expect a perfect match. As the QOM order increases, more and more non-zero effects match the signs of GWS eQTLs.

### 5 Omnigenes, higher-order QOM genes and PRS genes

We further characterise genes by the relative predictive power of individual models.

1. *QOM $\geq$ PRS genes* are where either of the QOMs $0^{th} - 3^{rd}$ outperforms or equals PRS.
2. *Omnigenes* are genes where the fraction of variance explained by QOM $3^{rd} \geq$ QOM $2^{nd} \geq$ QOM $1^{st} \geq$ QOM $0^{th}$ (*cis*) and QOM $3^{rd} \geq 0$

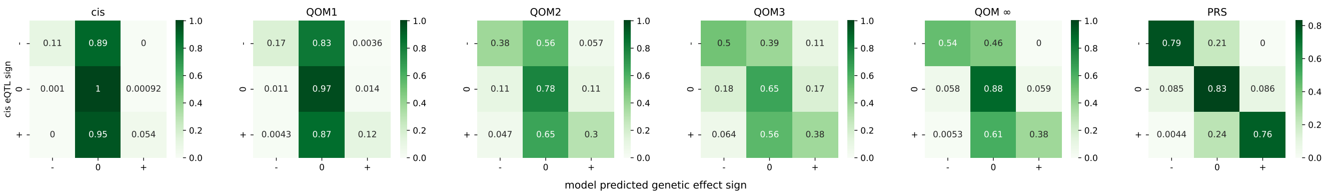


Figure SM4: **Comparison of effect directions between GWS eQTLs and the QOM.** The row label $i$ is the eQTL sign, column label $j$ denotes the sign of the genetic effect acc. to our model. The number in entry $(i, j)$ is the number of effects with sign GWS eQTL $i$ and model sign $j$, divided by the total number of GWS eQTLs of sign $i$ (normalised confusion matrix).

3. *Higher-order QOM genes* are genes $i$ where the performance of QOM bound (QOM $\infty$) is substantially larger than performance of the other QOMs, and at the same time close to PRS performance:

$$\max\left[R_i^2(\text{QOM } 0^{th} - 3^{rd})\right] < \epsilon_1 R_i^2(\text{QOM } \infty)$$
$$\&$$
$$R_i^2(\text{QOM } \infty) > \epsilon_2 R_i^2(\text{PRS})$$

4. *PRS genes* are genes where QOMs explain much less variance than PRS:

$$\max\left[R_i^2(\text{QOM } 0^{th} - 3^{rd}, \infty)\right] < \epsilon_3 R_i^2(\text{PRS}).$$

In Fig 2C we chose $\epsilon_1 = 0.6, \epsilon_2 = 0.8$ and $\epsilon_3 = 0.4$.

We note that each gene can be in one, multiple or none of the above categories. We can change the definitions of the above categories such as to account for uncertainty of estimated expressions (sub-sampling errorbars) – from "a gene fulfils a criterion whenever $a \geq b$" to "a gene fulfils a criterion whenever $A + \sigma_A \geq B - \sigma_B$", then the table from Fig 2C changes to:

| | |
|---|---|
| 183 | Transcription Factor genes |
| 143 (78%) | *QOM* $\geq$ *PRS* genes |
| 178 (97%) | *omnigenes* |
| 0 (0%) | *higher order QOM* genes |
| 5 (3%) | *PRS* genes |

Table 2: **Categories of genes when accounting for uncertainty of GE prediction.**

### 6 Regulatory network shuffles and controls

QOM is structured based on experimentally assembled GRNs. To test whether: *(i)* these GRNs carry predictive power, and *(ii)* whether QOM can be used as a test for the quality of network topology reconstruction, we train and evaluate QOM on randomised (shuffled) networks as a control. There are multiple ways to construct shuffled networks, conserving different topological and biological properties of the networks. We present three types of network shuffles and one control based on sampling markers at random.

**Topology-conserving shuffled networks** One way construct randomised networks by shuffling gene labels. These shuffles conserve the network topology, specifically, the number of links (number of free parameters) and their degree distributions. Results are shown in Fig 3 and Fig S8D-E.

If we did not conserve the degree distribution, a direct effect propagating through the shuffled network reaches, on average, *more* downstream genes than a direct effect in the experimentally-assembled GRN from Yeastract+ (which we refer to as the "true" GRN here, for simplicity). This is because the "true" GRN has a row-like topology with rows corresponding to out-degree hubs (master regulators) being nearly full. This structure is clearly not present in randomized networks.

We report that the "true" GRN is significantly more predictive than random controls, especially for high heritability genes. This is in part because low $h^2$ genes have lower in-degree than high $h^2$ genes (as suggested by Fig 2E). One possible explanation would be that a lower number of incoming regulatory links from upstream genes implies that the expression levels are less controlled (more noisy) and thus less heritable. In shuffled networks this correlation with heritability does not exist, and a QOM easily (over)fits these low $h^2$ genes.

**In-degree conserving shuffled networks** To test whether the improvement in the "true" network, compared to randomised networks, is or is not solely due to this heritability-in-degree correlation, we constructed a shuffle which only conserves the in-degree of each gene instead of the complete degree distribution, as schematised in Fig S8A. The "true" GRN QOMs perform significantly better than these specialized shuffles as well, on both high and low $h^2$ genes, as shown in Fig S8B-C.

In conclusion, the "true" network QOMs outperform topology-conserving and in-degree-conserving shuffles, the difference being larger for high $h^2$ genes. This improvement arises not solely due to the larger number of available free parameters (in-degree) for fitting these high $h^2$ genes, but rather because the "true" network more effectively captures the underlying regulatory interactions.

**Shuffled networks conserving functional grouping of genes** We also tested whether the actual transcriptional pathways in the GRN provide explanatory power, or whether the GRN only groups genes with similar functions. To address that, we divided the GRN into functional modules according to gene ontology (GO) annotations. For example, the TFs were assigned to one or more of $N = 12$ GO-Slim *biological function* categories. Based on that, we encoded each gene as a vector of zeros and ones in this N-dimensional space, and ran K-means clustering to cluster genes into $m$ modules (eg. $m = 5$, shown in S9A-C). The clustering was robust to initialization. Once the modules were defined, to get a shuffled network, we shuffled the gene labels only *within* the modules: "GO-shuffles" (schematised in S9A). As a control, we assigned the genes to these modules at random and, analogically, shuffled genes within the modules: "GO-controls". The results for $m = 5$ are shown in S9B,C. We repeated the QOM1 analysis also for $m = 9$ clusters obtaining qualitatively similar results (not shown). Altogether, these results show that the real GRN carries predictive power beyond grouping genes into functional modules.

**PRS-QOM controls** Another control we did, which we call the *PRS-QOM control*, was the following: QOM of order $k$ identifies a set of $N_i^{(k)}$ propagated effects (*trans*-eQTLs) for each gene $i$. For each order and gene, we randomly sample $N_i^{(k)}$ markers from the genome and ask how much variance these markers explain. If the GRN is predictive, the PRS on markers identified by the original QOM should perform better than PRS on $N_i^{(k)}$ randomly sampled markers. More precisely, we sample the markers such that the number of independent LD blocks for each gene, and the distribution of number of markers per LD block is the same as for the original QOM. To make a fair comparison, we exclude the corresponding *cis* regions for every gene (since the direct effects are not determined by the network), thus putting only the *trans*-eQTLs identified by the GRN to test. Results from this PRS-QOM control analysis shown in S9D confirm that the PRS-QOM based on "true" network outperforms corresponding controls at all orders, serving as a further proof that the experimentally assembled GRN carries predictive power.

Last, we note that it is not trivial to estimate the expected performance of QOMs trained on randomised networks. Suppose that the "true" network consisted solely of causal regulatory links, causing the expression levels of genes to be strongly correlated (as Fig 4 shows). When we shuffle this causal network, non-causal links appear, which, however, still capture the low-dimensional, correlated pattern of variability between genes. If expression levels of gene $A$ and gene $B$ are correlated because they are both causally regulated by $C$, the "true" network would contain links $C \to A$ and $C \to B$, but the shuffled network could well contain a link $A \to B$ or $B \to A$, which is also statistically predictive. Therefore, it is not surprising that shuffled networks do perform relatively well. What is important, however, is that the "true", experimentally assembled GRN performs significantly better than properly randomized controls.

### 7 QOM using network models based on "Expression Evidence"

We replicate the main results for *Expression evidence* based GRN assembled by the Yeastract+ consortium [1] (SI Figure fig. S2). As specified in SI Appendix Sec. 1, *Expression evidence* based GRN is defined using co-expression patterns, which means such GRN may by design contain links which are not causal, and correspond to only *correlation* between expression levels of a regulator and a target.

Therefore, *Expression evidence* based GRN is larger ($G = 217$) and less sparse (6062 regulatory links; 12.9% of entries of $\mathbf{B}$ are potentially nonzero) as compared to the *DNA binding evidence* GRN ($G = 183; 2864$ regulatory links; 8% sparsity). Because there are more regulatory links in GE evidence networks, QOM models are less sparse (especially higher-order QOMs, as shown in Fig. fig. S3), automatically allowing the less-constrained higher-order GE-evidence-based QOMs to explain more heritability than the more-constrained DNA-binding-evidence-based QOMs. In contrast, while *cis*, $1^{st}$ *order* QOM, and *PRS* performances are nearly identical between the two networks.

Qualitatively, both DNA-binding- and GE-evidence-based QOMs show increase in performance with higher orders of effect propagation, in line with the omnigenic hypothesis (SI Fig. fig. S2A, E). Both types of evidence lead to significantly higher heritability of in-degree hubs as compared to non-hub genes, but the difference is smaller for GE-evidence-based QOMs (SI Fig. fig. S2C); similarly, both sets of QOMs predict expression of the in-degree hubs better than of non-hub genes, while there is no significant difference between out-degree hubs and non-hubs (SI Fig. fig. S2D).

While GE-evidence-based QOMs have more degrees of freedom than DNA-binding-evidence-based QOMs and thus in general higher prediction performance, shuffled controls reveal a more nuanced picture (SI Fig. fig. S2F-H). Here, GE-evidence-based QOM of the $2^{nd}$ and $3^{rd}$ *order* outperform network shuffles for high $h^2$ genes, similarly for DNA-binding-evidence-based QOMs in the main paper. However, $1^st$ *order* GE-evidence-based QOM does *not* outperform the corresponding shuffles, unline the DNA-binding-evidence-based QOM. There are two possible reasons for this, of which the first effect is likely the dominant one that naturally explains the discrepancy.

We reason about the dominant effect as follows. By nature of detection, as explained in Yeastract+ database, GE evidence is composed of "non-causal correlation" links and real "causal" links. Many of the non-causal links will arise precisely due to effect propagation: if the causal interactions were $A \to B$ and $B \to C$, GE-evidence-based GRN would likely include also the link $A \to C$ link (since all three genes, A, B, and C would be co-regulated). At higher orders (but not at first order), correlation and causation may blend together: $A \to C$ is also causal, not only correlative, at $2^{nd}$ and $3^{rd}$ order effect propagation. This could explain why GE-evidence-based QOMs outperform shuffles at $2^{nd}$ and 3rd order but not at $1^{st}$ order. At $1^{st}$ order, causal and non-causal interactions are distinct, and while GE-evidence-based QOM has more links (and thus more parameters to fit) than DNA-binding-evidence-based QOM and thus would have the potential to outperform it – it doesn't, it has nearly indistinguishable performance – its shuffles do have more free parameters and thus make the experimentally-assembled network performance insignificantly better than shuffles, unlike in the DNA-binding-evidence-based QOMs presented in the main text. In other words, GE-evidence-based QOMs of $1^{st}$ order contain causal and spurious, non-causal links; non-causal links do not improve the fit on the experimentally-assembled network, but do improve it on network shuffles, making the performance of the experimentally-assembled network insignificantly different from shuffles. If this explanation holds, it provides further argument that comparison with network shuffles can be used as a test of causal network topology.

The second possible effect relevant for the interpretation of our results is as follows. Since GE-evidence is based on observed correlation in GE change due to a perturbation, this detection may be biased towards larger effects; therefore causal/ correlation links that only cause small changes in GE could be missing. An equivalent bias would likely be smaller for DNA-binding-evidence-based GRNs. Since populations with varying frequencies of polymorphic sites were used to assemble the GRNs, effects that would be small in data-collection environments could be large in our population. This could lead to the GE-evidence-based GRN to miss some crucial links present in DNA-binding-evidence-based GRN, thus decreasing the power of the corresponding QOMs, which would mostly be visible at the $1^{st}$ *order.*

### 8 Gene expression covariance

We compute covariance matrices, $\mathbf{C}$, for expression values predicted by each model, and ask how well the covariance seen in the experiment, $\mathbf{C}_{\text{exp}}$, is captured by the models based on known gene regulatory network structure. We first do element-wise comparisons of the model-predicted covariance matrices (specifically, its off-diagonal elements) to the data covariance matrix, by computing squared Pearson correlation ($R^2$, fraction of variance explained), as shown in Fig 4C.

**Significant covariance.** We evaluate which pairs of genes are significantly correlated by comparing to a null distribution of covariance elements, computed as follows. GE values for each individual (rows of $\mathbf{Y}_{\text{exp}}$) are independently shuffled and covariance matrix is computed for such shuffled $\mathbf{Y}_{\text{exp}}$. This is repeated 50 times, and 2.5%

and 97.5% quantiles are taken to be the significance thresholds for entries of $\mathbf{C}_{\text{exp}}$ (grey lines in Fig 4B). Choosing different quantiles only affects the results of the Extended QOM (Fig 5), and that only very slightly (for $q = 0.01$, the slopes in Fig 5D remain unchanged and $R^2$ changes by at most 0.02).

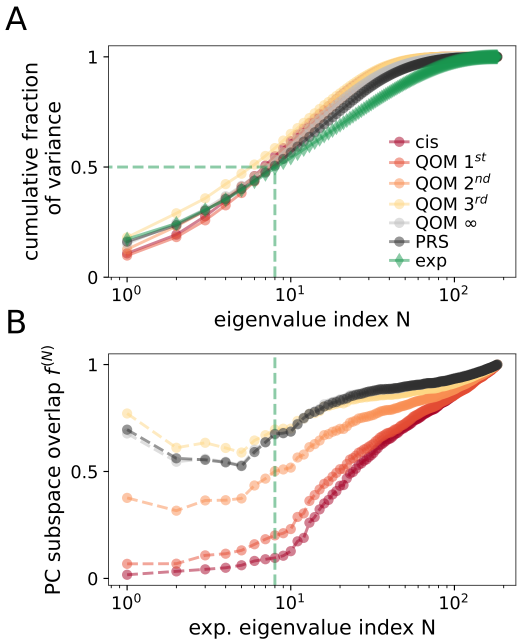


Figure SM5: **Measure of overlap of PC subspaces between various models and data. (A)** Cumulative sum of ordered eigenvalues, for each model and experimental data, same as Fig. 4D. **(B)** Ratio of the fraction of variance explained by first N PCs of $\mathbf{Y}_{\text{exp}}$ and the fraction of variance explained by experimental PCs, $f^{(N)}$, defined in (M16). 50% of variance in data is explained by the first 8 eigenvectors (dashed green line).

**PCA, models vs. experiment overlap.** To asses whether the structure of the covariance is lower-dimensional, and whether the sub-spaces of dominant variability align between the experimental data and the models, we use principal component analysis (PCA) on the $\mathbf{C}$ for all models and $\mathbf{C}_{\text{exp}}$ computed on testval set of individuals, and evaluate the overlap between these subspaces as follows. For each model prediction $\mathbf{Y}$ and individual $i$ from the testval set, we compute the squared length of the expression vector $y_i$ in the subspace spanned by the first (top) $N$ eigenvectors of $\mathbf{Y}_{\text{exp}}$: $|y_i^{(N)}|^2$, average across individuals to get $\langle |y^{(N)}|^2 \rangle$, and normalize by $\langle |y|^2 \rangle$, to get the fraction of data vector variance that lies within the subspace of the first $N$ PCs of experimental data. This is plotted in Fig 4E for each model separately. For data, this procedure is simply equivalent to the cumulative sum of the eigenvalues (fraction of variance of data explained), plotted in Fig 4E. For models, this projecting procedure can be understood as a modified PCA, where we compute the variance in model prediction explained by *experimental* PCs, and not by model's own PCs.

To define overlap of PC subspaces $f^{(N)}$ shown in Fig. SM5, for each $N$, we divide this average length $\langle |y^{(N)}| \rangle$ by the average length of the projection of the experimental expression vector:

$$f^{(N)} = \frac{\langle |y^{(N)}|^2 \rangle / \langle |y|^2 \rangle}{\langle |y_{\text{exp}}^{(N)}|^2 \rangle / \langle |y_{\text{exp}}|^2 \rangle} \tag{16}$$

### 9 Extended QOM: propagation of non-transcriptional effects

Regulatory pathways propagate genetic transcriptional as well as genetic non-transcriptional and non-genetic - environmental - effects (we refer to the latter two as non-transcriptional effects or noise - effectively, in the simple QOM, both genetic non-transcriptional and environmental effects enter as residuals or "noise", since they do not originate from *cis*-eQTLs.). Some of this non-transcriptional variability will propagate through the regulatory network and contribute, in a correlated fashion, to the variability of downstream genes. The basic expectation is therefore that non-transcriptional variability *should not* be independent across genes. Moreover, some of this co-variability should be predictable precisely by the same statistical model (and putative underlying mechanism) that we use to propagate genetic effects. To verify this hypotheses, we formulate an extended version of the Quantitative omnigenic model:

$$\tilde{\mathbf{Y}} = \mathbf{XD} + \mathbf{N} + \sum_{k=1}^{K} \alpha_k (\mathbf{XD} + \mathbf{N})\mathbf{B}^k + \tilde{\mathbf{E}} \tag{17}$$

where $\tilde{\mathbf{Y}}$ are expression levels, $\mathbf{X}$ , $\mathbf{D}$ and $\mathbf{B}$ are defined as in the genetic QOM (2), $\tilde{\mathbf{E}}$ are residuals and $\mathbf{N}$ is the "noise" or non-transcriptional effects (non-transcriptional genetic and environmental) matrix, $\mathbf{N} \sim \mathcal{N}(0, \mathbf{\Sigma})$. We assume $\mathbf{\Sigma}$ to be a diagonal matrix with "noise variances of each gene", $\sigma_i^2$, on the diagonals. These variances can represent sources of variability due to the different environmental / internal state of each cross population. Specifically, since the experiments are done at the population level, these "noise" terms are unlikely to be due to direct biochemical, "intrinsic" stochasticity within individual cells, and are more likely due to what in systems biology literature is referred to as "extrinsic" noise sources. $\mathbf{N}$ is further assumed to be independent of both the *cis* genetic effects $\mathbf{XD}$ and the extended QOM residuals $\tilde{\mathbf{E}}$.

Our goal is to explain some of the covariance unexplained by the genetic QOM by propagation of random non-transcriptional effects $\mathbf{N}$ through the same GRN $\mathbf{B}$ fitted by the genetic QOM. To do that, let's imagine $\mathbf{N}$ and $\tilde{\mathbf{E}}$ are known. Then we can write the experimental gene expression levels as

$$\mathbf{Y}_{\text{exp}} = \mathbf{XD} + \mathbf{N}(\mathbb{1} + \tilde{\mathbf{B}}_K) + \tilde{\mathbf{E}} \tag{18}$$

and the empirical gene-gene covariance averaged across individuals $\mathbf{C}_{\text{exp}} \in \mathbb{R}^{G \times G}$, where $G = 183$ is the number of genes, as:

$$\begin{aligned}
\mathbf{C}_{\text{exp}} &= \frac{1}{N}\mathbf{Y}_{\text{exp}}^T \mathbf{Y}_{\text{exp}} && (19)\\
&= \frac{1}{N}\left[\left(\mathbb{1} + \tilde{\mathbf{B}}_K\right)^T \mathbf{D}^T\mathbf{X}^T\mathbf{XD}\left(\mathbb{1} + \tilde{\mathbf{B}}_K\right) + \left(\mathbb{1} + \tilde{\mathbf{B}}_K\right)^T \mathbf{N}^T\mathbf{N}\left(\mathbb{1} + \tilde{\mathbf{B}}_K\right) + \tilde{\mathbf{E}}^T\tilde{\mathbf{E}}\right] + && (20)\\
&+ \frac{1}{N}\left\{(\mathbb{1} + \tilde{\mathbf{B}}_K)^T\mathbf{D}^T\mathbf{X}^T\mathbf{N}(\mathbb{1} + \tilde{\mathbf{B}}_K) + (\mathbb{1} + \tilde{\mathbf{B}}_K)^T\mathbf{N}^T\tilde{\mathbf{E}} + \tilde{\mathbf{E}}^T\mathbf{XD}(\mathbb{1} + \tilde{\mathbf{B}}_K)\right\} + && (21)\\
&+ \frac{1}{N}\left\{(\mathbb{1} + \tilde{\mathbf{B}}_K)^T\mathbf{D}^T\mathbf{X}^T\mathbf{N}(\mathbb{1} + \tilde{\mathbf{B}}_K) + (\mathbb{1} + \tilde{\mathbf{B}}_K)^T\mathbf{N}^T\tilde{\mathbf{E}} + \tilde{\mathbf{E}}^T\mathbf{XD}(\mathbb{1} + \tilde{\mathbf{B}}_K)\right\}^T && (22)
\end{aligned}$$

In the limit of large population size $N$ and under the above stated assumptions, the cross terms $\{.\}$ go to zero.[1] We assume that our sample size ($N = 800$ training individuals) is sufficient to consider the cross terms $\{.\}$ negligible compared to the other terms. Then we obtain a simplified equation $\mathbf{C}_{\text{exp}} = \mathbf{C}_{\text{G}} + \mathbf{C}_{\text{N}} + \mathbf{C}_{\tilde{\text{E}}}$, where

$$\mathbf{C}_{\text{G}} = \frac{1}{N}\left(\mathbb{1} + \tilde{\mathbf{B}}_K\right)^T \mathbf{D}^T\mathbf{X}^T\mathbf{XD}\left(\mathbb{1} + \tilde{\mathbf{B}}_K\right) \tag{23}$$

$$\mathbf{C}_{\text{N}} = \left(\mathbb{1} + \tilde{\mathbf{B}}_K\right)^T \mathbf{\Sigma}\left(\mathbb{1} + \tilde{\mathbf{B}}_K\right). \tag{24}$$

$\mathbf{C}_{\text{G}}$ is the empirical genetic transcriptional, $\mathbf{C}_{\text{N}}$ the non-transcriptional covariances and $\mathbf{C}_{\tilde{\text{E}}}$ is the covariance that remains unexplained by the extended QOM.

However, in reality, $\mathbf{N}$ and $\tilde{\mathbf{E}}$ are not known. Nevertheless, we can infer the postulated *variances* of the propagating "noise" component $\mathbf{N}$, which will be the unknown vector $\vec{x}$:

$$\vec{x} = \text{diag}\,(\mathbf{\Sigma}) = (\sigma_1^2, \sigma_2^2, ..., \sigma_G^2). \tag{25}$$

We infer $\vec{x}$ from the covariance unexplained by the transcriptional effects alone $\mathbf{C}_{\text{exp}} - \mathbf{C}_{\text{G}}$, which is known: $\mathbf{C}_{\text{exp}}$ is computed from measured expressions as in (M19) and $\mathbf{C}_{\text{G}}$ is computed from predicted expressions (from (M23) given $\tilde{\mathbf{B}}_K$ and $\mathbf{D}$ fitted by the genetic QOM). Thus, we formulate a minimization problem for the unknown $\vec{x}$ with the associated loss function:

$$\text{Loss} = \sum_{i,j}\left[\mathbf{C}_{\text{exp}} - \mathbf{C}_{\text{G}} - \left(\mathbb{1} + \tilde{\mathbf{B}}_K\right)^T \text{diag}(\vec{x})\left(\mathbb{1} + \tilde{\mathbf{B}}_K\right)\right]_{ij}^2. \tag{26}$$

[1] the genetic transcriptional and noise effects are orthogonal in $N \to \infty$ limit, therefore $\mathbf{D}^T\mathbf{X}^T\mathbf{N} \to (0)$, $\mathbf{N}^\mathbf{T}\tilde{\mathbf{E}} \to \mathbb{0}$ and $\tilde{\mathbf{E}}^T\mathbf{XD} \to \mathbb{0}$

This loss is quadratic in $\vec{x}$, and we use quadratic programming (QP) (python library `cvxopt` implementation) to solve the optimization problem. With the following definitions: $\mathbf{Z} = \mathbf{C}_{\text{exp}} - \mathbf{C}_{\text{G}}$ and $M_j = \tilde{\mathbf{B}}_K^T * \tilde{\mathbf{B}}_{(:,j)} = \tilde{\mathbf{B}}_K^T * (\tilde{b}_{K1j}, \tilde{b}_{K2j}, ..., \tilde{b}_{KGj})^T$, where $*$ denotes element-wise multiplication, and

$$\mathbf{P} = 0.5 \sum_j M_j^T M_j \tag{27}$$

$$q = -\sum_j M_j^T \mathbf{Z}_{(:,j)}, \tag{28}$$

the loss can be re-written in the canonical QP form:

$$\text{Loss} = \sum_j (M_j \vec{x} - \mathbf{Z}_{(:,j)})^T (M_j \vec{x} - \mathbf{Z}_{(:,j)}) = \vec{x}^T \mathbf{P} \vec{x} + \vec{x}^T \vec{q}. \tag{29}$$

Since elements of $\vec{x}$ represent variances, we set the constraint $x_i \geq 0$, and the upper bound is given by the total non-transcriptional variance for each gene, $\vec{x} \leq \text{diag}(\mathbf{Z})$.

We apply this method to the $3^{rd}$ *order* model ($K = 3$).

**Direct vs. propagated transcriptional and non-transcriptional variance ($\xi, \eta$).** Since GRNs propagate genetic transcriptional as well as non-transcriptional effects, we asked whether genes whose expression is mostly influenced by *trans* genetic effect (effects propagated through transcriptional regulatory pathways) instead of direct *cis* effects display the same characteristic for non-transcriptional (genetic and environmental) effects. We define and compare two ratios:

$$\vec{\xi} = \frac{\text{“trans”}}{\text{“cis+trans (total genetic transcriptional)”}} = 1 - \frac{\text{diag}(\mathbf{D}^T \mathbf{X}^T \mathbf{X} \mathbf{D})}{\text{diag}(\mathbf{C}_{\text{G}})}. \tag{30}$$

$$\vec{\eta} = \frac{\text{“propagated non-transcriptional”}}{\text{“total non-transcriptional”}} = 1 - \frac{\text{diag}(\mathbf{\Sigma})}{\text{diag}(\mathbf{C}_{\text{N}})} = 1 - \frac{\vec{x}}{\text{diag}(\mathbf{C}_{\text{N}})},$$

where $\mathbf{D}$ are the direct effects inferred by the cis model (M2), $\mathbf{N}$ are non-transcriptional effects as stated in (M17), $\mathbf{C}_{\text{G}}$ and $\mathbf{C}_{\text{N}}$ are transcriptional genetic and non-transcriptional covariance matrices defined by (M23) and (M24), respectively and $\vec{x}$ are the inferred “noise” variances as defined by (M25). In Fig 5C we observe that the genes with high ratio of propagated transcriptional to total transcriptional variance ($\xi \geq 0.5$) are enriched in genes with high ratio of propagated non-transcriptional to total non-transcriptional variance ($\eta$). We call these genes “sensitive”: sensitive to upstream regulators rather than direct effects. Expression of these “sensitive” genes is mostly influenced by propagation through transcriptional regulatory pathways.

**Analytical variance decomposition.** To determine whether the extended QOM framework can explain existence of ”sensitive” genes, i.e. the observed connection between $\xi$ and $\eta$, we look at the analytical variance decomposition of the extended QOM (M17)). Covariance of gene $j$ can be written down as:

$$\text{Cov}(\vec{y}_j) = \text{E}\left[\vec{y}_j \otimes \vec{y}_j^T\right] \tag{31}$$

$$\text{Cov}(\vec{y}_j) = \text{E}\left[\left[(\mathbf{XD} + \mathbf{N})(\mathbb{1} + \tilde{\mathbf{B}})\right]_j \otimes \left[(\mathbf{XD} + \mathbf{N})(\mathbb{1} + \tilde{\mathbf{B}})\right]_j^T\right] \tag{32}$$

$$\text{Cov}(\vec{y}_j) = \text{E}\left[(\mathbf{X}\vec{d}_j + \vec{n}_j + \mathbf{XD}\vec{\tilde{b}}_j + \mathbf{N}\vec{\tilde{b}}_j) \otimes (\mathbf{X}\vec{d}_j + \vec{n}_j + \mathbf{XD}\vec{\tilde{b}}_j + \mathbf{N}\vec{\tilde{b}}_j)^T\right] \tag{33}$$

where $\vec{y}_j$ is the $(1 \times N)$ vector of expression levels for gene $j$ for $N$ individuals. Similarly, $\vec{d}_j$, $\vec{n}_j$ and $\vec{\tilde{b}}_j$ is $j^{th}$ columns of the direct effects matrix $\mathbf{D}$ , non-transcriptional/ noise effects matrix $\mathbf{N}$ and propagated effects matrix $\tilde{\mathbf{B}}$ defined in (M5), respectively. $\vec{d}_j \sim \mathcal{N}(0, \mathbb{1}\sigma_{dj}^2)$, $\vec{n}_j \sim \mathcal{N}(0, \mathbb{1}\sigma_{nj}^2)$ and $\tilde{b}_{ij} \sim \mathcal{N}(0, \sigma_{b_{ij}}^2)$.

We further assume the following: transcriptional and non-transcriptional effects are independent, and direct effects are independent of propagated effects, since our GRN $\mathbf{B}$ has zeroes on diagonals. Higher order loops exist ($\tilde{\mathbf{B}}$ has nonzero diagonals, in general) but in our case this results in negligible (and insignificant, according to threshold in to Fig 3B) covariance. Therefore, expectations of all the cross-terms in (M33) are zero, and we get a simplified expression for the $N \times N$ covariance matrix:

$$\mathrm{Cov}(\vec{y}_j) = \mathrm{Cov}(\mathbf{X}\vec{d}_j) + \mathrm{Cov}(\vec{n}_j) + \mathrm{Cov}(\mathbf{XD}\vec{\tilde{b}}_j) + \mathrm{Cov}(\mathbf{N}\vec{\tilde{b}}_j) \tag{34}$$

$$\mathrm{Cov}(\vec{y}_j) = \mathbf{XX^T}\sigma^2_{d_j} + \sigma^2_{e_j} + \mathbf{XX^T}\sum_{k=1}^{G}\sigma^2_{d_i}\sigma^2_{\tilde{b}_{ij}} + \sum_{k=1}^{G}\sigma^2_{e_k}\sigma^2_{\tilde{b}_{kj}} \tag{35}$$

To connect the above to (M30), we compute the average variance for gene $j$ across individuals $\frac{1}{N}\mathrm{Tr}\left[\mathrm{Cov}(\vec{y}_j)\right]$, and get expression for $\xi$ and $\eta$:

$$\xi_j = 1 - \frac{\sigma^2_{d_j}}{\sum_{k=1}^{G}\sigma^2_{d_k}\sigma^2_{\tilde{b}_{kj}}} \tag{36}$$

$$\eta_j = 1 - \frac{\sigma^2_{e_j}}{\sum_{k=1}^{G}\sigma^2_{e_k}\sigma^2_{\tilde{b}_{kj}}}. \tag{37}$$

While $\xi$ and $\eta$ are not strictly equal, there is a potential explanation for why $\xi$ and $\eta$ could be correlated. When gene $j$ has a large number of regulators, we can replace the sums in (M36) by expectations (under the assumption that $\frac{\sigma^2_{d_k}}{\sigma^2_{d_j}}$ and $\frac{\sigma^2_{e_k}}{\sigma^2_{e_j}}$ are independent of $\sigma^2_{\tilde{b}_{kj}}$). If, additionally, the strengths of the non-transcriptional noise effects and genetic transcriptional effects on a given gene are correlated, e.g. when the origin of noise is biophysically connected to the binding of TFs onto the promoter, we could assume that $\frac{\sigma^2_{d_k}}{\sigma^2_{d_j}} \propto \frac{\sigma^2_{e_k}}{\sigma^2_{e_j}}$. In that case, $\xi_j$ is expected to be proportional to $\eta_j$.

**Comparing covariance matrices.** We compare similarity of the gene-gene covariance of predicted expressions of the $3^{rd}$ *order* QOM model and extended QOM model of Eq. (17), $\mathbf{C}$, to the covariance of experimentally measured expressions, $\mathbf{C}_{\mathrm{exp}}$. The key idea is that including the propagation of non-transcriptional effects/ "noise" in the extended QOM model can explain significantly larger fraction of experimentally estimated covariance in gene expression. We compare two measures evaluated over the significant pairwise correlations of the data covariance matrix, $\mathbf{C}_{\mathrm{exp}}$, collected into a vector, $\vec{c}_{\mathrm{exp}}$ (and the corresponding model values collected into $\vec{c}$), with significance determined as in Section 8. On these selected covariance elements we evaluate:

1. Fraction-of-variance explained, using Pearson correlation $R^2 = \left[r(\vec{c}, \vec{c}_{\mathrm{exp}})\right]^2$.
2. The slope, $\beta$, of the linear regression, $\vec{c} = \beta\vec{c}_{\mathrm{exp}}$. While the $R^2$ values are similar for simple QOM and extended QOM, the simple QOM has a much smaller slope ($\beta_G$) than the extended model ($\beta_{G+N}$), as shown in Fig 5D. The extended model thus not only correlates with, but also explains the absolute magnitude of the significant non-transcriptional covariance elements.

We note that part of the gene-gene covariance explained by the Extended QOM could be attributed to non-transcriptional regulatory hotspots. Genetic effects picked up by PRS overlap significantly more regulatory hotspots identified by Albert et al [1] than expected by chance, and majority of these hotspots (65/95) do not overlap a transcription factor. Therefore, we hypothesise that effects of these non-TF hotspots (which contribute to the gap between QOM and PRS) could be partially captured by the Extended QOM.

### 10 Benchmark on synthetic data

To verify our methodology, we generated matrices $\mathbf{D}_{\mathrm{sim}}$ and $\mathbf{B}_{\mathrm{sim}}$, from which we generated synthetic gene expression data using the SEM-derived form of the model (main text Eq (2) where $K \to \infty$ or solution of Eq (3)):

$$\mathbf{Y}_{\mathrm{sim}} = (\mathbf{X}\mathbf{D}_{\mathrm{sim}} + \mathbf{N})\left(\mathbb{1} - \mathbf{B}_{\mathrm{sim}}\right)^{-1} + \mathbf{E} \tag{38}$$

where $\mathbf{E}_{ij} \sim \mathbb{N}(0, \sigma^2_{\mathrm{int}})$ and $\mathbf{N}_{ij} \sim \mathbb{N}(0, \sigma^2_{\mathrm{ext}})$ are intrinsic and extrinsic (propagated) noise.

We choose $\mathbf{D}_{\mathrm{sim}}$ to have the same sparsity structure as fitted $\mathbf{D}$: $d_{\mathrm{sim}\,ij} = 0$ if polymorphism $i$ does not affect gene $j$ according to inferred *cis* model (M2). Otherwise, $d_{\mathrm{sim}\,ij} = d$ where $d$ is drawn from the distribution of inferred direct effects. Similarly, we chose $\mathbf{B}_{\text{sim}}$ to have the same structure as $\mathbf{B}$ in QOMs (taken from Yeastract+), $b_{\text{sim}ij} = 0$ when gene $i$ is not a direct regulator of gene $j$. Otherwise, $b_{\text{sim}ij} = \kappa b$, where $b$ are drawn from the distribution of fitted regulatory weights inferred by $3^{rd}$ *order* QOM (shown in Fig S4C bottom). $\kappa = 0.35, \sigma_{\text{int}} = 0.4, \sigma_{\text{ext}} = 0.15$ were chosen such that the distributions of simulated expressions levels $\mathbf{Y}_{\text{sim}}$ and gene-gene covariances are as close to their empirical counterparts as possible. The comparison is shown in Fig S1A, B.

Fig S1C-F evaluates how precisely can the QOM models (M1) and (M7) and the PRS model (M9) reproduce the ground truth genetic effects and regulatory strengths.

First, in Fig S1C we compare the average per gene fraction-of-variance explained by the ground truth matrices $\mathbf{B}$ and $\mathbf{D}$ vs and the inferred matrices $\mathbf{B}$ and $\mathbf{D}$ , $\mathbf{G}_{\text{QOM}}$ and $\mathbf{G}$ . We define the ground truth predicted expressions as follows:

$$\text{QOM 0-3: } \hat{\mathbf{Y}}_{\text{GT}} = \sum_{k=0}^{K} \mathbf{X}\mathbf{D}_{\text{sim}}\mathbf{B}_{\text{sim}}^{k} \tag{39}$$

$$\text{QOM bound and PRS: } \hat{\mathbf{Y}}_{\text{GT}} = \mathbf{X}\mathbf{D}_{\text{sim}}(\mathbb{1} - \mathbf{B}_{\text{sim}})^{-1} \tag{40}$$

and the inferred model predictions (analogous to fitting real data):

$$\text{QOM 0-3: } \hat{\mathbf{Y}} = \sum_{k=0}^{K} \mathbf{X}\mathbf{D}\mathbf{B}^{k} \tag{41}$$

$$\text{QOM bound: } \hat{\mathbf{Y}} = \mathbf{X}\mathbf{G}_{\text{QOM}} \tag{42}$$

$$\text{PRS: } \hat{\mathbf{Y}} = \mathbf{X}\mathbf{G}. \tag{43}$$

We define the performances as average per gene fraction-of-variance explained $\langle R_i^2 \rangle$, where $R_i^2 = r^2(\hat{y}_{i\text{GT}}, y_{i\text{sim}})$ for predictions using ground truth, and $R_i^2 = r^2(\hat{y}_i, y_{i\text{sim}})$ for inferred models.

As anticipated, with the ground truth matrices the performance rises with QOM order, while for the inferred effects, performance peaks at QOM3. Here QOM bound and PRS are overparametrised and hence can perform worse than QOM3. This trend differs from fitting real expression data, as depicted in, for example, Fig 1C. The reason may be that in the synthetic case defined by (M38), all heritable variance comes from direct effects propagating through the known GRN. However, non-transcriptional effects contribute to variance in real expression levels, which can be captured by the less restricted QOM bound and PRS models. Further, inferred models can capture some of the noise variance propagated through the same GRN, and can thus outperform predictions on ground truth matrices, as observed for lower order QOMs in Fig S1C.

Fig S1(D-F) shows the detailed inferred vs ground truth comparisons. (D) compares individual matrix entries of $\mathbf{D}$ and $\mathbf{B}$ , $\mathbf{G}_{\text{QOM}}$ and $\mathbf{G}$ (E) compares LD-corrected genetic effects and finally (F) compares expression levels. The simulation demonstrates that direct effects and regulatory weights can be effectively inferred, with a higher precision observed as the QOM order increases, as can be seen in (D). Accounting for LD leads to improved agreement, as shown in (E).

### 11 QOM applied to a natural yeast population

We replicated the key parts of our analysis on a pan-population of yeast, sampled from various geographical and ecological niches forming a representative sample of the species. The pan-population was genotyped by Peter et al (2018) [**Peter2018**] and profiled for gene expression by Caudal et al (2024) [**Caudal2024**]. The individuals fall into 29 clades, ranging in size from one to 354 strains. The genetic distances between strains are relatively within clades and large between clades (Fig S10F). Nevertheless, given the large divergence of clades, their gene expression patterns were relatively similar [**Caudal2024**]. The MAF and GE distributions for both the pan-population and the cross are shown in Fig S10G,H.

**Genotypes.** In our analysis we used 943 strains for which both genotypes (1 754 867 markers) and gene expression measurements (6445 genes) were available. We ran our analysis on 83 794 sites with MAF $> 5\%$ used in GWAS by Ref [**Peter2018**] plus the markers we defined as direct effects (explained below in sec. Direct effects). The strains varied in ploidy. We encoded their genotypes according the original study as 0/1/2 for 0, 1 and 2 or more copies of an alternative allele and normalized by alternative allele frequency $p_j$ as $X_{ij} = x_{ij} - 2p_j$. We filled missing values with average genotype in each clade whenever possible, and with the pan-population average otherwise. We chose

750 individuals at random for training and used the remaining 193 individuals divided into 93 and 100 in each fold, for evaluation (as defined in sec. 1).

**LD pattern.** The yeast pan-population consists of strains which are evolutionary relatively divergent from one another (Fig. S10E), and each clade consist of relatively small number of closely related individuals - the lineages coalesce far back in the past. This means that most MAF $> 5\%$ polymorphisms are fixed in one or multiple clades and in most cases we do not observe within-population genetic variation. This implies large LD between variants all over the genome, with effectively no decay with physical distance from the markers (Fig. S10B). This LD pattern presents a challenge in identifying (causal) variants linked with gene expression differences between strains, since there is a large number of highly correlated variants all over the genome which could explain these between-clade expression differences. In other words, most variants are associated only with a small number of genetic backgrounds and environments, lacking the resolution of the Albert et al eQTL-mapping study population we use for our main analysis (Fig. S10A).

Looking at individuals clades, the LD pattern can decay with distance from the marker (Fig. S10C), similar to the cross (Fig. S10A). However, the clades separately contain too few individuals to conduct clade-specific GWAS, and most clade-specific polymorphisms have a MAF $< 5\%$ and thus were filtered out in the main analysis.

**Direct effects.** We defined the direct effects matrix $\mathbf{D}$ based on the GWS *cis*-eQTLs detected by the original analysis [**Caudal2024**]. Whenever the GWS *cis*-eQTL variant was not available in the genotypes, we picked the closest marker. We included markers with MAF $< 5\%$ among direct effects, since only 61 *cis*-eQTLs were present in the original GWAS genotypes by Ref [**Peter2018**]. Due to the non-trivial LD structure described above, we included only *cis*-eQTL markers as direct effects (not a whole genomic window), thus making a conservative estimate.[2] The direct effects matrix is shown in Fig S10E. Out of 183 TFs, only 24 had a *cis*-eQTL (we call these genes *cis*-genes) and out of these 24 *cis*-genes, only 15 had a non-zero out-degree (can propagate direct effects). This is much less than 93/110 *cis*-genes with non-zero out degree in the cross. Moreover, the average out-degree of *cis-genes* is lower than of a typical TF (Fig S11B). Therefore, as expected, the power to detect eQTLs in this natural pan-population is much lower than in the cross. We further observe that these GWS *cis*-eQTLs are more linked with one another than a random, equally large set of markers randomly sampled from the genome (Fig S10D).

**Gene expression levels.** Gene expression measurements were available for 180 out of 183 regulators according to Yeastract+ DNA binding evidence GRN. The three missing genes were `YIR017C, YKR034W` and `YLR013W` whose expression levels we replaced by that of genes with most similar expressions according to the SPELL database. These were `YNL277W, YJR152W` and `YOL015W`, respectively. We further standardised the gene expression levels defined as $y_{ij} = \log_2(1 + \text{TPM}_j)$ (TPM = transcripts per million) as $Y_{ij} = \frac{y_{ij} - \overline{y_j}}{\text{std}(y_j)}$.

**QOM results.** The main results of QOM performance on the yeast pan-population are shown in Fig S11. We show that the QOM explains variance in expression via propagation through transcriptional regulatory pathways, with higher orders explaining more variance. In particular, despite only 15 *cis*-genes which can propagate direct effects to other TFs downstream, $1^{st}$ order QOM predicts gene expression of 30 additional TFs with no *cis*-eQTLs ($p < 0.05$), doubling the variance explained by the $0^{th}$ *order* QOM. Expression of further 61 genes ($p < 0.05$) is predicted by $2^{nd}$ *order* QOM as compared to $1^{st}$ *order* QOM, again increasing average-per-gene variance explained twofold. Adding $3^{rd}$ order effect propagation further increases model performance, reaching 2/3rds of that of QOM bound and 42% of the amount of GE variance explained by unrestricted PRS. The absolute amount of variance in GE explained is lower than for the cross, which is expected due to the lower statistical power as discussed above, and a small number of detected GWS *cis*-eQTLs (direct effects). This also explained the relatively larger gaps between $3^{rd}$ *order* QOM, QOM bound and PRS.

As in the main analysis, QOM identifies various types of genes (Fig S11C). There are relatively more *PRS genes* (14%) and less *QOM¿PRS genes* (3%) and *omnigenes* (25%) as compared to the cross. This is expected due to the substantially smaller fraction of markers the QOM uses to fit gene expression levels, resulting in overall smaller

[2] We repeated the analysis by defining the direct effects matrix as GWS *cis*-eQTLs and markers within 250bp window upstream and downstream of each eQTL (typical $\text{LD}_{1/2}$ distance for this dataset is 500bp [**Peter2018**]), increasing the number of direct effect from 265 to 2266. Model performances remained similar, however, we could not reproduce the significant difference between shuffled and "true" network. We suspect this is because not all markers in 500bp windows are in LD with the *cis*-eQTL and include other markers in LD with *trans*-eQTLs in the rest of the genome, thus confounding the analysis.

number of genes the QOM can predict at every order. Nevertheless, for the pathways which can be identified, we observe relatively high consistency between QOMs of various orders (Fig S11E).

The two example genes from Fig S10I are highlighted in Fig S11E. `YKL109W`, which is is differentially expressed between domesticated and wild clades, does not have a GWS *cis*-eQTL and the expression is explained solely by effect propagation through the GRN. Each subsequent QOM order explains additional variance, and the identified pathways are highly consistent. In contrast, most variance in expression of `YDR520C` is driven by direct effects present in all individuals of the Ecuadorean clade. Trans-acting variants identified by higher order models and PRS further contribute to the low expression level of `YDR520C` in this clade. We hypothesise there could be more such clade-specific signal which did not reach genome-wide significance in the smaller clades, in part explaining the small number of GWS *cis*-eQTLs.

To conclude, QOM successfully utilizes prior pathway information in predicting gene expression of TFs even in natural populations where statistical power is limited. Since QOM predicts gene expression via effect propagation through transcriptional GRN, it could, in principle, suggest causal *trans*-eQTLs despite the high correlation of causal variants with a large fraction of the genome.

**QOM shuffles and controls.** Fig S11D shows that QOMs trained on the "true" network outperform QOMs trained on shuffled networks, suggesting that the GRNs are predictive and can identify causal interactions also in natural populations with low MAF and complex LD structure (Fig S10B-G). In particular, since the 24 *cis*-genes have a lower out-degree than a typical gene (S11B), we performed out-degree conserving shuffles (analogous to S8A but shuffling only the *column* labels) to ensure a fair comparison. We report significant improvement of "true" QOM in the $1^{st}$ and $2^{nd}$ order, as compared to the corresponding shuffles.

Further, we ran the PRS-QOM control described in SI sec. 6. Since the LD does not have a simple block structure, it is not trivial to correct for and therefore we ran a naive control without any LD correction. We don't report a significantly higher performance of the "true" network (not shown). We believe that one of the reasons is that the *cis*-eQTLs are more linked to each other than a random set of markers (S10D). Therefore, a random set of markers in PRS-QOM control is on average more independent, resulting in higher coverage of the genome than the *cis*-eQTLs in the "true" PRS-QOM. In order to perform a fair PRS-QOM control on this pan-population dataset, a less trivial control correcting for non-block-like LD needs to be designed. This was beyond the scope of our analysis and we leave this for future explorations.

### Supplementary figures

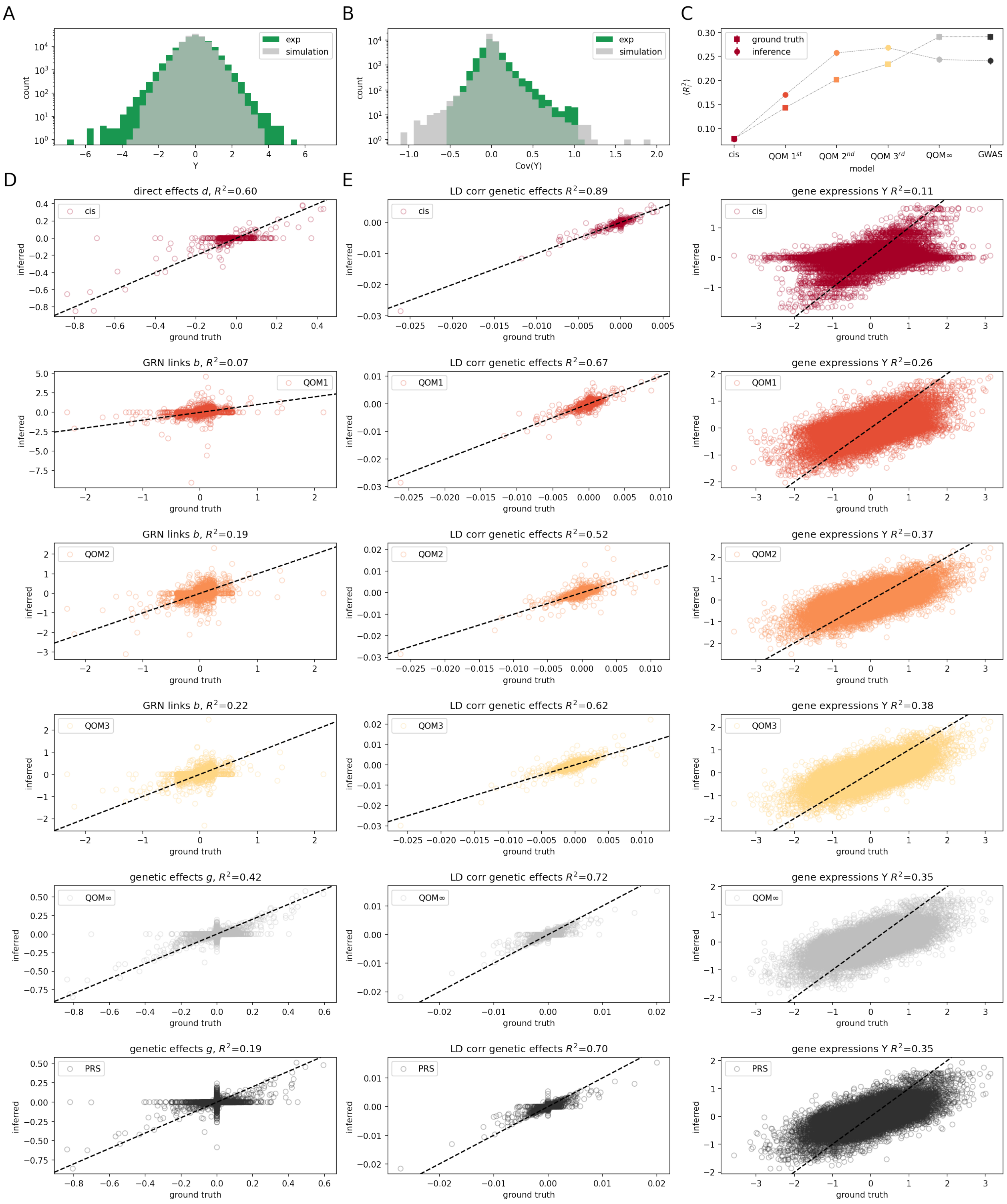


Figure S1: **Synthetic data.** **(A)** distribution of synthetic and real gene expression levels $\mathbf{Y}_{\text{exp}}$ and $\mathbf{Y}_{\text{sim}}$. **(B)** Distribution of gene-gene covariance, for synthetic and real data. **(C-E)** Comparison of inference and ground truth for each model. In the models, **B** and **D** were substituted with inferred and ground truth matrices, as Eq. (M39)-(M43) specify. **(C)** Average per gene fraction-of-variance explained. Errorbars corresponds to the resamping standard error across folds and are within size of the markers. **(D)** *A-priori* nonzero entries of matrices **D** for the *cis* model, **B** for QOM models, and all genetic effects $\mathbf{G}_{QOM}$ and **G** for QOM bound and PRS model. **(E)** Genetic effects matrices corrected for LD. **(F)** Comparison of predicted and generated expression levels $\hat{\mathbf{Y}}$ vs $\mathbf{Y}_{\text{sim}}$.

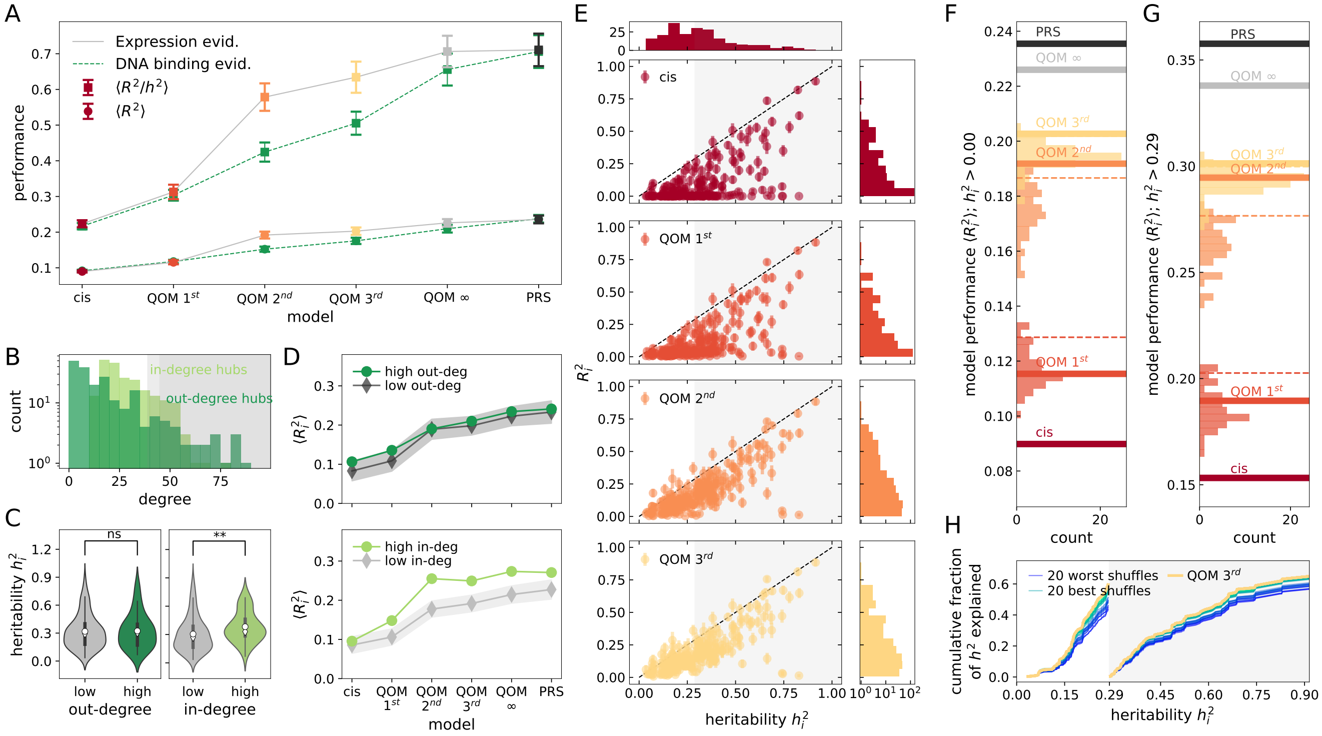


Figure S2: **Key results GE-evidence-based QOMs.** **(A)** Measures of model performance for individual models based on GE-evidence-based GRN (grey, color) in comparison to models trained on DNA-binding-evidence-based GRN (green). Plotted is average per gene fraction-of-variance explained ($\langle R_i^2 \rangle$, circle) and mean fraction of GE heritability explained ($\langle R_i^2/h_i^2 \rangle$, square). Model performance errorbars are SD across 50 resamplings of 112 individuals from the evaluation set (SI Appendix Sec. 1). Higher-order GE-evidence-based QOMs outperform their DNA-binding-evidence-based counterparts. **(B)** Distribution of in- and out-degree of genes. We call the top 10% (15%) of genes in- (out-) degree hubs, with in-(out-) degree above 39 (45). **(C)** Distribution of $h^2$ for hubs vs non-hub genes. No significant difference between genes with high ($\langle h^2 \rangle = 0.323$) and low ($\langle h^2 \rangle = 0.317 \pm 0.028$, 100 resamplings) out-degree ($p > 0.5$, U-test). In-degree hubs have significantly higher $h^2$ ($\langle h^2 \rangle = 0.38$) than non-hub genes ($\langle h^2 \rangle = 0.31 \pm 0.03$, 100 resamplings; $p < 0.003$, U-test). White circle signifies the mean, diamond the median. **(D)** $\langle R_i^2 \rangle$ of different models for hubs vs non-hubs. Shaded areas denote standard error of the mean across 100 samples of $H$ non-hub genes, where $H$ is the number of hub genes ($H = 39$ in- and $H = 34$ out-degree hub genes). Unlike out-degree hubs, in-degree hubs have significantly higher $\langle R_i^2 \rangle$ than non-hub genes, the difference being highest for QOM1-3. **(E)** Per-gene variance explained, $R_i^2$, as a function each gene's heritability ($h_i^2$), estimated by Albert *et al* [1]. Errorbars are standard deviations across samples (SI Appendix Sec. 1). Marginal distributions of $h_i^2$ and $R_i^2$ shown on the sides. **(F, G)** Models based on experimentally-assembled GE-evidence-based GRN (solid lines) and shuffled GRNs (distributions). Shuffling procedure replicates that of main text Fig. 3A. Plotted is average-per-gene variance explained, $\langle R_i^2 \rangle$, evaluated on (F) all genes and (G) genes with above median heritability $h_i^2 > 0.29$. Number of shuffled networks for $1^{st}$, $2^{nd}$ and $3^{rd}$ *order* QOMs is 51, 52 and 77, respectively. Dashed lines show 5% significance cuts in the matched shuffled distributions. **(H)** Cumulative sum of per gene fraction-of-variance explained, normalised by the sum of heritability across genes, $\sum_{i \in S} R_i^2 / \sum_{i \in S} h_i^2$, as a function of $h_i^2$, plotted separately for a set of genes $S$ with $h^2$ below and above the median (0.29). Results shown for the $3^{rd}$ *order* QOM: thick yellow line corresponds to GE-evidence-based GRN, thin green (blue) lines to the 20 best (worst) performing shuffled GRNs.

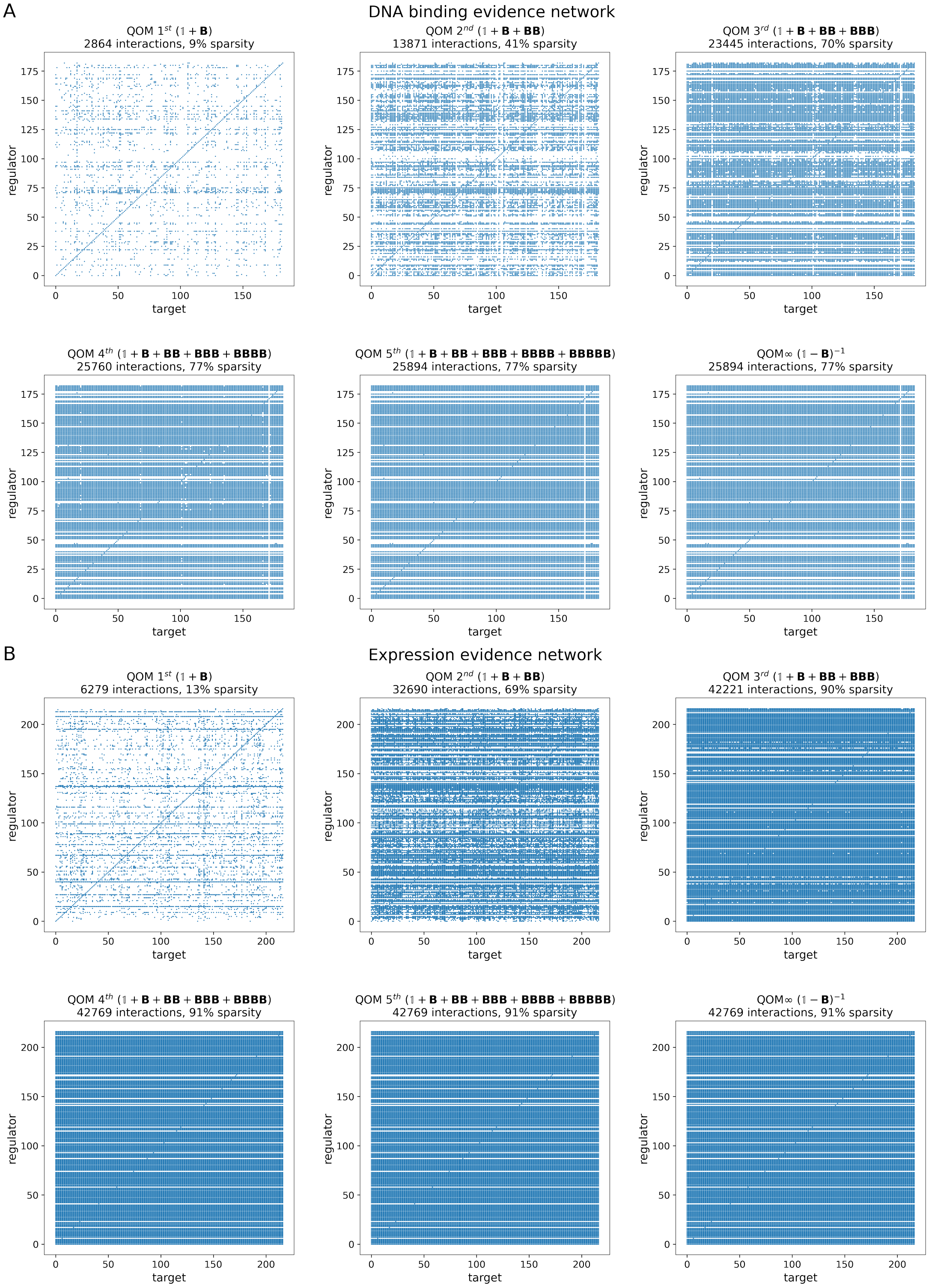


Figure S3: **Patterns of effect propagation through the GRN in QOMs of various orders for (A) DNA binding evidence and (B) Expression evidence GRN.** Each plot shows the structure of propagated effects network $\mathbb{1}+\tilde{\mathbf{B}}_K = \sum_{k=0}^{K} \mathbf{B}^k$ (as defined in Eq (M5)). Blue = observed, white = unobserved regulatory interactions. Shown are all possible direct and indirect regulator-target interactions in QOM of order $K$ allowed by regulatory evidence in the Yeastract+ database. The number of regulatory interactions saturates at $5^{th}$ order in (A) and $4^{th}$ order in (B) and matches the *structure* of $(\mathbb{1}-\mathbf{B})^{-1}$ and $\tilde{\mathbf{B}}_\infty$, defined by Eq (M7) and Eq (M5). In both (A) and (B), the $3^{rd}$ is less than 2.5 thousand indirect interactions ($< 10\%$) away from saturation. Expression evidence networks are less sparse as compared to DNA evidence networks, especially for higher order QOMs.

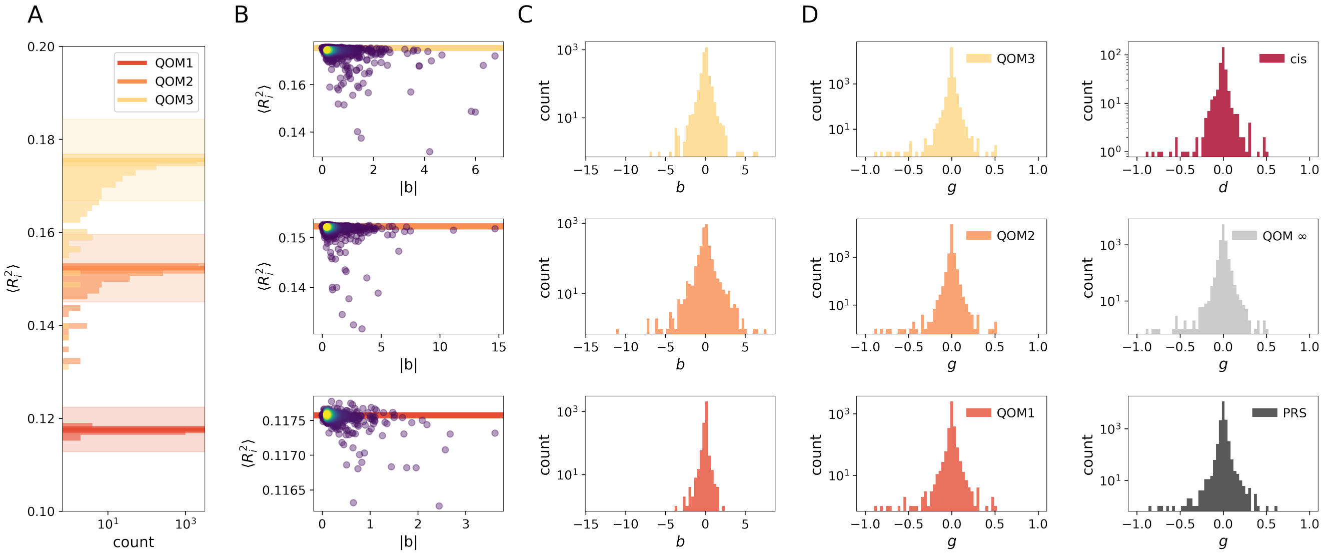


Figure S4: **Impact of inferred regulatory links, their size distribution, and distribution of propagated effect magnitudes.** **(A)** Distribution of performances of QOMs after removing one link at a time without re-fitting the model. Thick lines show the performance of corresponding full QOMs (no links removed), shaded areas denote the standard deviation and standard error of the mean of the full QOM performance, obtained by resampling. Most link removals lead to a slight decrease in performance. **(B)** Decrease in performance (vertical axis) is significantly correlated with the magnitude of the link (horizontal axis). $b$ denotes the regulatory link weights: entries of matrix $\mathbf{B}$ as defined by (2) or (M3) and (M4). **(C)** Distributions of fitted regulatory links $b$ and **(D)** distribution of total genetic effects for each model. $d$ are individual *cis* effects: entries of $\mathbf{D}$ defined by (M2). $g$ are entries of matrices $\mathbf{G}_K, \mathbf{G}_{\text{QOM}}$ and $\mathbf{G}$ defined by (M6), (M7) and (M9), respectively.

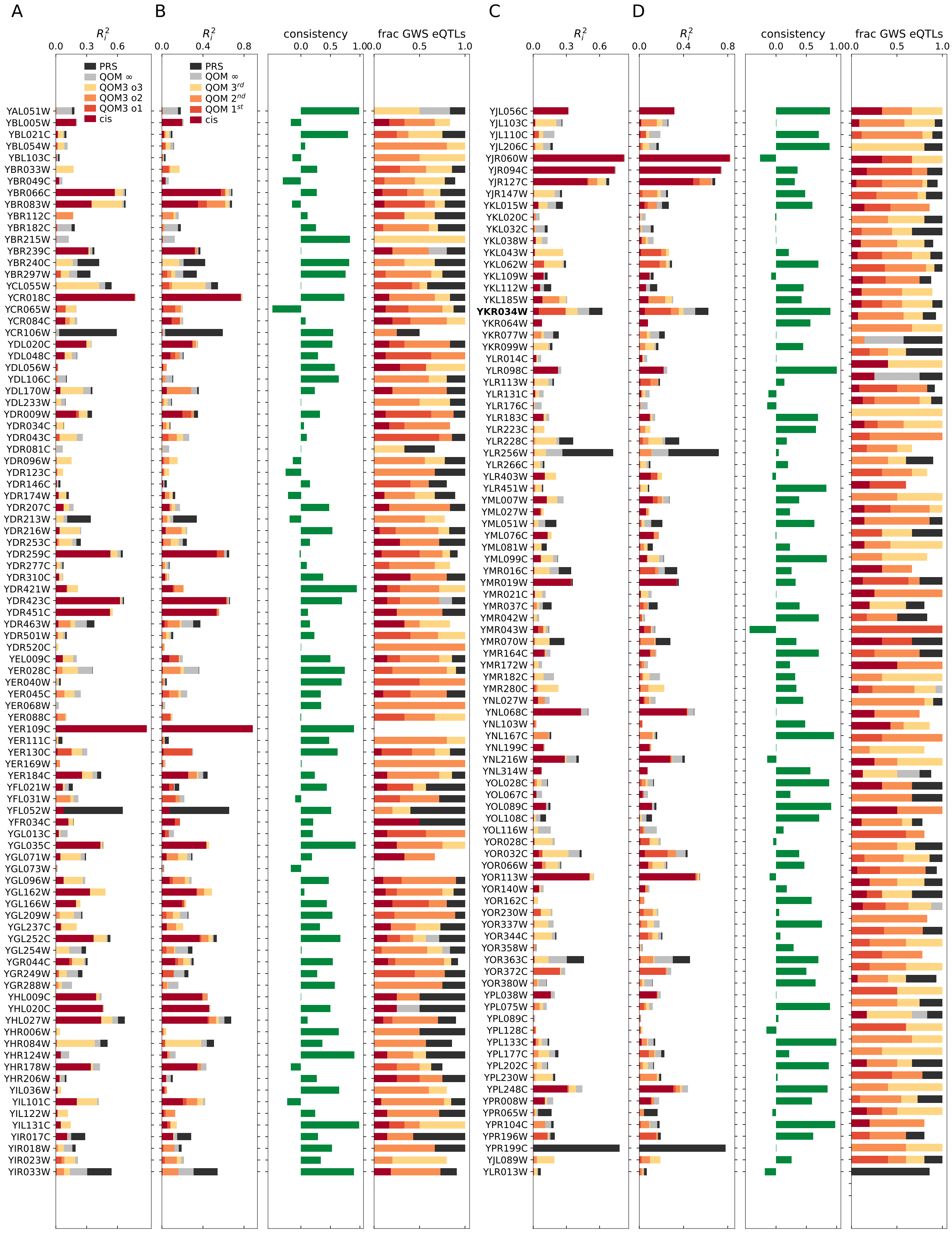


Figure S5: **Breakdown of order-by-order performance for all genes.** **(A, C):** Contribution of individual orders *within* a single fit of $3^{rd}$ *order* QOM ($K = 3$). Cumulative genetic contributions up to order $k, k \leq K$ are defined as $\mathbf{G}_k = \mathbf{D} \sum_{i=0}^{k} \mathbf{B}^i$ (analogously to (M6)). Performance metrics defined as in main text Fig. 2D. **(B, D)** Order-by-order performance evaluated by *separate* fits of QOM models of different orders, $K$. This is a complete (all genes) version of Fig. 2D of the main text, where only selected genes were shown.

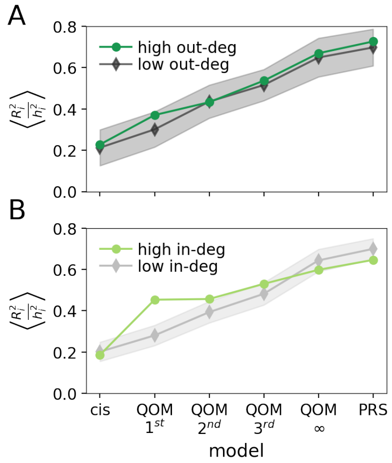


Figure S6: **Average per gene fraction of heritable variance explained by each model for hub and non-hub genes**. Analog to Fig 2E of the main text. **(A)** No significant difference between out-degree hubs vs non-hubs. **(B)** QOM 1-3 perform better in in-degree hubs than on non-hubs. Shaded area is a standard deviation of mean fraction of heritable variance explained over 20 randomly drawn sets of N=27 in (A) and N=45 (B) non-hub genes.

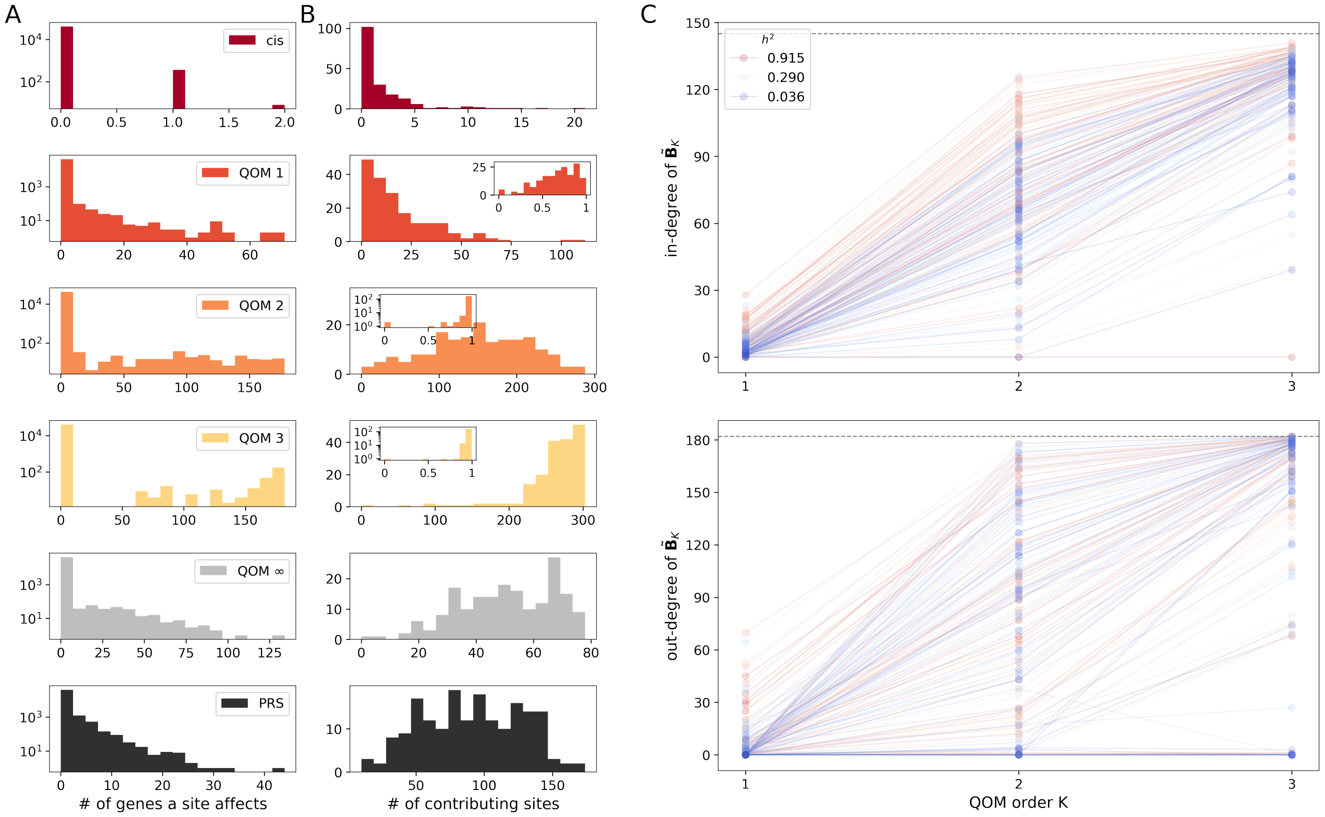


Figure S7: **Analysis of polygenicity.** **(A)** Distribution across sites of the number of genes each of the 42 052 polymorphic sites affects, for each model (rows). With increasing QOM order, most *cis*-eQTLs end up additionally contributing as *trans*-eQTLs to the expression level of most of the genes. The plotted distributions thus tend towards a bimodality, which differs from the PRS-type model. **(B)** Distribution across genes of the number of polymorphic sites (out of 42 052) affecting each of the 183 genes. For cis and QOM 1-3, these can only be sites which are a *cis*-eQTL to a regulator. Insets show the distribution of the ratio between the actual number of sites contributing to a given gene and the maximal number of sites that could contribute to that gene given the GRN topology. Maximum number of contributing sites is close to saturated already in QOM $3^{rd}$, implying that three steps of effect propagation seem to explain most of the polygenicity resulting from transcriptional regulation. The pattern of polygenicity revealed by QOM $3^{rd}$ substantially differs from what an unstructured PRS model suggests. While PRS relies heavily on regularisation, QOMs are constrained mainly by the GRN structure. **(C)** Propagated effects network topology defines the pattern of polygenicity. Plots show the total in- and out- degree for each gene given by the structure of the propagated effects network $\tilde{\mathbf{B}}_K$ (M5) at order $K$, $K \in \{1, 2, 3\}$. Data points corresponding to one gene are connected by a line and colored by gene's heritability. Dashed lines denote the maximum in-degree (145) and out-degree (182) given the structure of $\tilde{\mathbf{B}}_\infty$ defined by (M5). The in-degree of most genes saturates fast, after two or three steps of propagation, explaining the prevalent polygenicity. Higher heritability genes tend to have higher in-degree, in agreement with Fig.2E. Higher heritability genes have higher out-degree at $K = 1$, since only regulatory links from regulators with a *cis*-eQTL can be estimated from $1^{st}$ *order* QOM and these genes are typically more heritable than genes with no *cis*-eQTL. Out-degree for most regulators also saturates at $2^{nd}$ or $3^{rd}$ order.

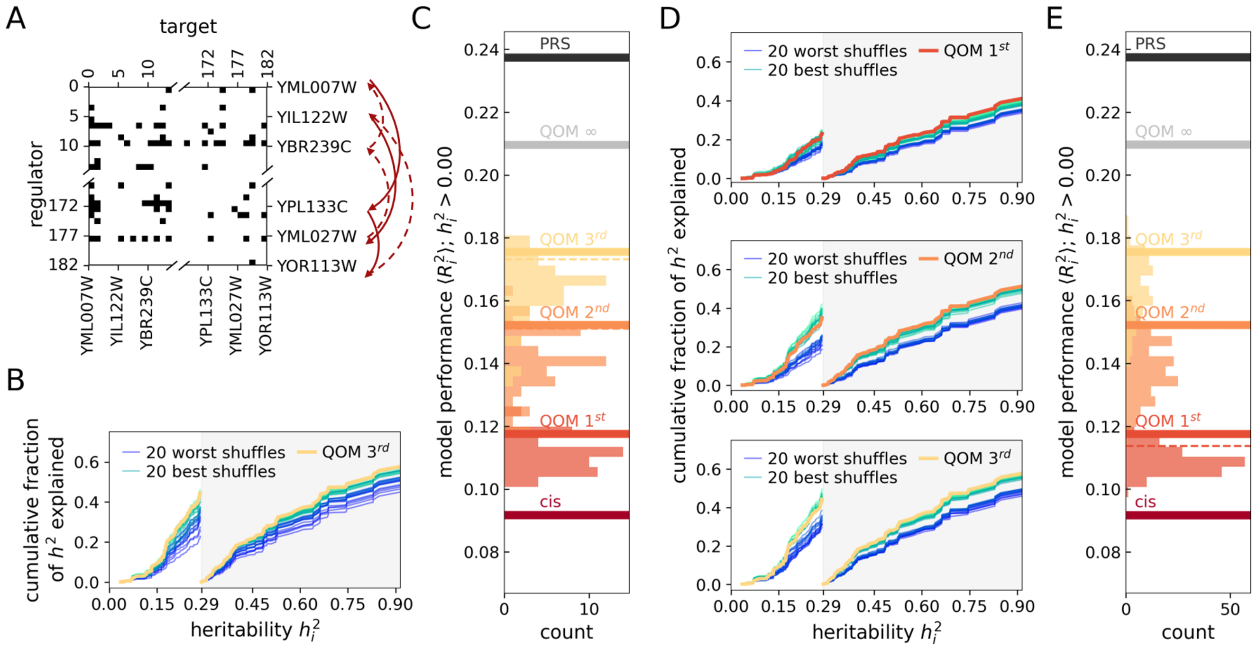


Figure S8: **Experimentally assembled GRN outperform both in-degree conserving (A-C) and topology-conserving (D-E) shuffled networks.** **(A)** Shuffling of networks by re-naming regulators (i.e. shuffling only row labels) conserves the in-degree of all genes, and thus conserves the correlation between heritability and in-degree. **(B, D)** Cumulative distribution function of $R_i^2$ of genes as a function of their $h_i^2$. Thick lines correspond to QOM performance given true GRN, thin green (blue) lines to the 20 best (worst) performing shuffled GRNs. The true network outperforms most shuffled networks, especially for high $h^2$ genes for topology-conserving shuffles (D), and for all genes for in-degree conserving shuffles (B). **(C, E)** Models based on true GRN (lines) and shuffled GRNs (distributions). Shown is average-per-gene $\langle R_i^2 \rangle$, evaluated on genes in the whole heritability range. Dashed lines denote $p = 0.1$ significance threshold (where not visible, dashed and thick lines merge). A set of 52 in-degree conserving shuffled networks was used to fit all QOM models in (C). The number of topology-conserving shuffled networks in (E) is 111, 168 and 94 for QOM for QOM $1^{st}$, $2^{nd}$ and $3^{rd}$ order, respectively.

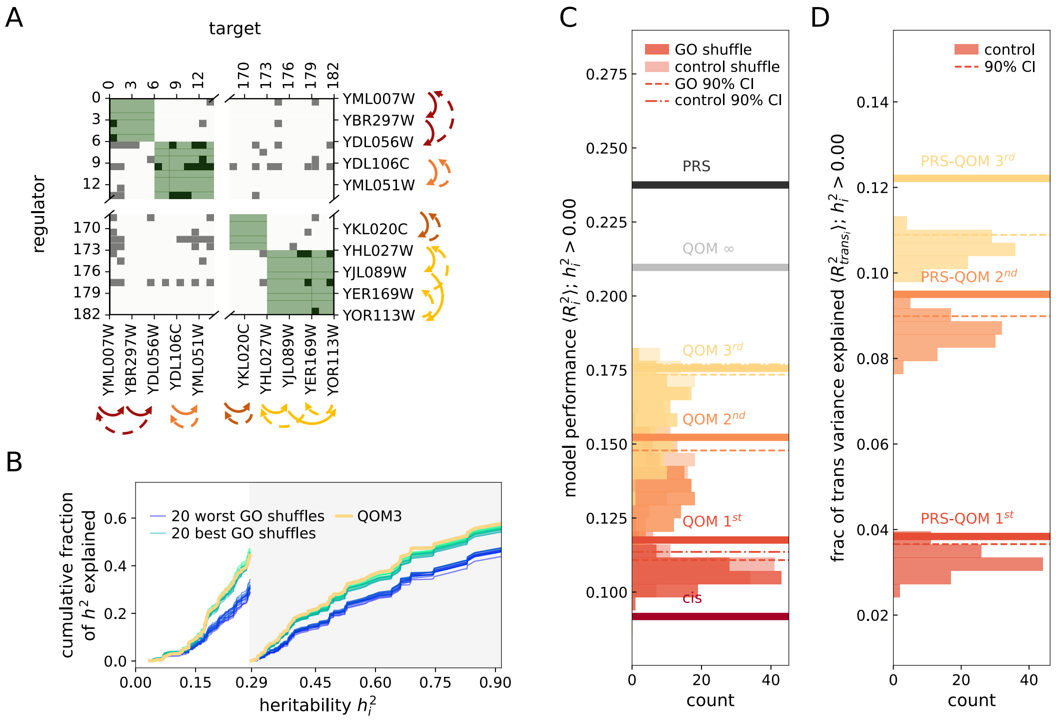


Figure S9: **Experimentally assembled GRN outperforms controls: is predictive beyond functional grouping of genes (A-C) and network-implied trans-eQTLs explain more variance than a random set of markers (D). (A)** Shuffling of networks by re-labelling genes within each functional module. Modules are defined by clustering genes according to their GO biological function annotations ("GO-shuffles") or into random modules of the same sizes ("controls"). This shuffle conserves the connectivity within a functional module, while shuffling the transcriptional regulatory links. **(B)** Cumulative distribution function of $R_i^2$ of genes as a function of their $h_i^2$. Thick lines correspond to QOM performance given true GRN, thin green (blue) lines to the 20 best (worst) performing GO-shuffles. The "true" network outperforms most shuffled networks, especially for high $h^2$ genes. **(C)** Models based on true GRN (lines), GO-shuffles (dark distributions) and control shuffles (light distributions). Shown is the average-per-gene $\langle R_i^2 \rangle$, evaluated on genes in the whole heritability range. Dashed lines denote $p = 0.1$ significance threshold. A set of X networks conserving functional grouping and the same number of controls was used to fit all QOM models. "True" network outperforms both GO-shuffles and control shuffles. **(D)** Fraction of per-gene *trans* variance explained by PRS-QOM. PRS on *trans*-eQTLs identified by QOM on "true" networks (lines), vs PRS on a random set of markers (excluding cis regions for each gene) (distributions) and $p = 0.1$ significance threshold (dashed line). True network-based markers explain more variance than a random set of markers.

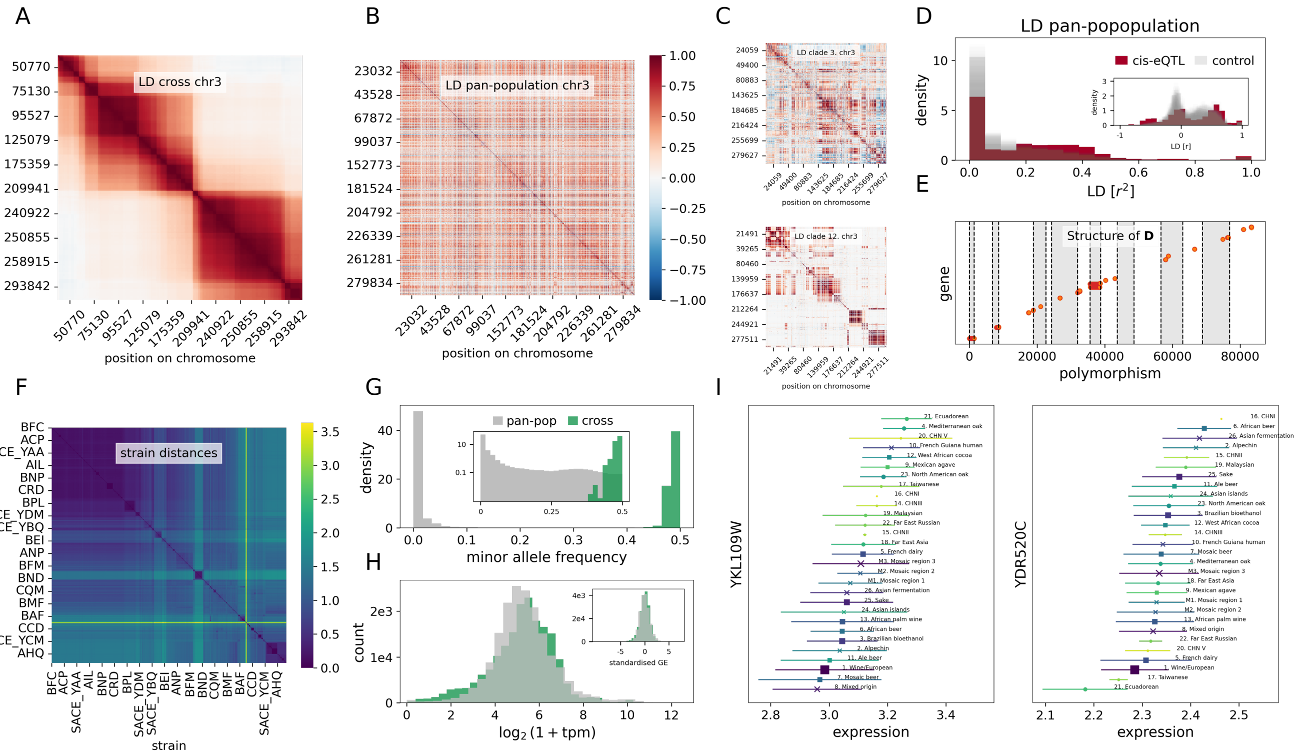


Figure S10: **Comparison of pan-population vs cross.** **(A-C)** LD on chromosome 3 in (A) yeast cross population, (B) the whole pan-population and (C) for two clades from the pan-population (10.French Guiana human and 12. West African cocoa) plotted separately. Color indicates LD defined as the Pearson correlation between each pair of sites, from -1 (dark blue) to +1 (dark red). **(D)** LD between 61 GWS *cis*-eQTLs with MAF > 5% (red) vs LD between a random set of 61 markers with MAF > 5% (100 draws) plotted in grey. *cis*-eQTLs are more tightly linked than markers in the rest of the genome. **(E)** Structure of the direct effects matrix **D** based on GWS *cis*-eQTLs reported by [**Caudal2024**] (and the closest variant to a GWS cis-eQTL if the *cis*-eQTL was not genotyped) . Out of the 663 potential direct effects (orange dots), 265 were chosen by L1 regularisation in the *cis* model (red dots), implying that in QOMs, the variance in expression of all genes of interest is explained solely by these 265 sites. **(F)** SNP distances for each pair of strains, in percentage of non-identical bases. Heterozygous differences were half-weighted compared to the homozygous differences [**Caudal2024**]. Distances are small within clades (¡1%) while distances between clades are large, up to 3.5%. **(G)** Minor allele frequency (MAF) distribution of pan-population (grey) vs in the cross (green), density in linear (main fig) and log scale (inset). **(H)** Distribution of gene expressions measured in $log_2(1 + \mathrm{TPM})$ for pan-population (grey) and cross (green). Standardised expression plotted in the inset. GE distributions are similar between the two datasets. **(I)** Gene expressions by clade for two example *cis*-genes. The marker and errorbar indicate mean and standard deviation of expression in each clade. The size and color of the marker correspond to clade size: from 1 individual (CHIN I, yellow) to 354 (Wine/European, dark blue). Marker shape signifies clade domestication: square: domesticated; circle: wild; cross: unassigned. Differential expression (DE) of `YKL109W` is driven by more commonly segregating variants associated with domestication, while DE of `YDR250C` is driven by strong cis-eQTL present in the Ecuadorean clade.

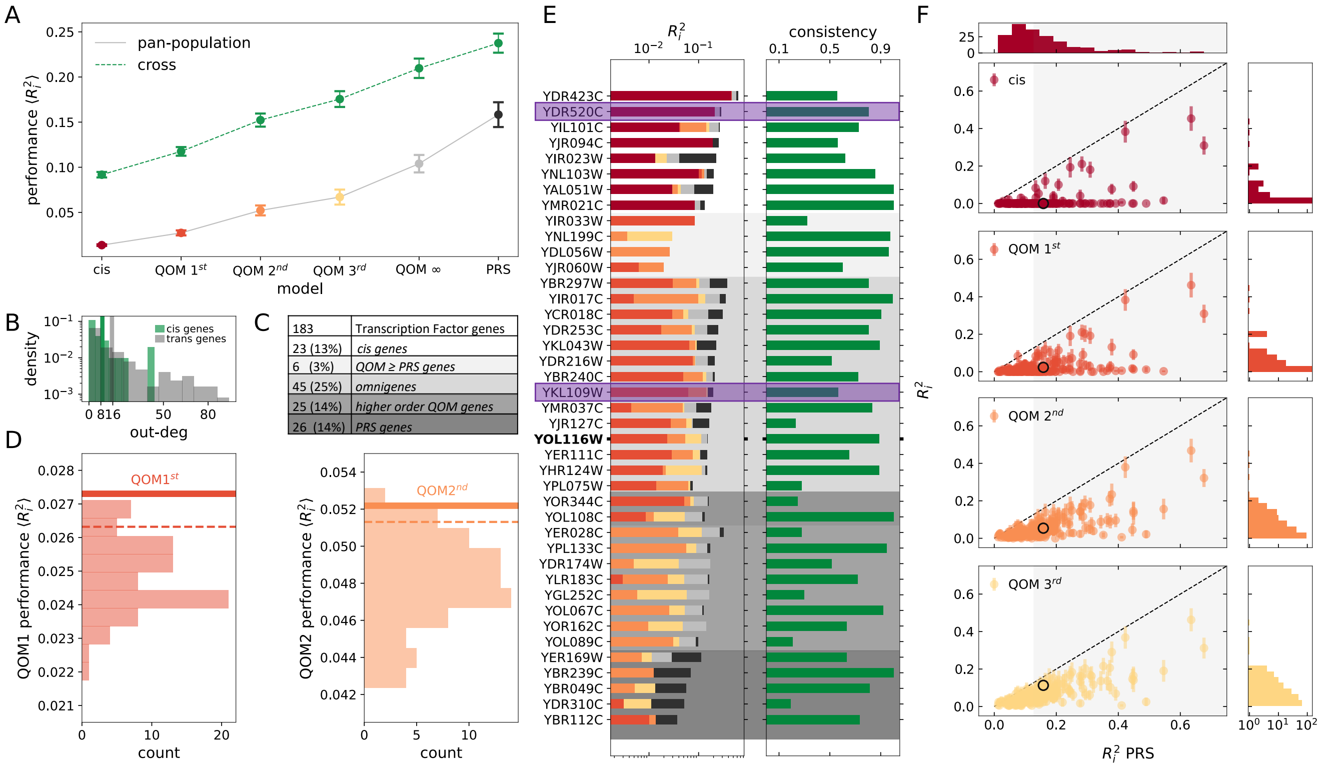


Figure S11: **Key results from natural yeast pan-population.** **(A)** Measures of model performance for individual models based on pan-population (grey, color) in comparison to models trained on the yeast cross (green), in both cases on DNA-binding-evidence-based GRN. Plotted is average per gene fraction-of-variance explained ($\langle R_i^2 \rangle$). Model performance errorbars are SD across 50 resamplings of individuals from the evaluation set (SI Appendix Sec. 1). **(B)** Out-degree distributions of genes with a GWS *cis*-eQTL (green), and the remaining (*trans*) genes with no direct effects (grey). *Cis* genes have a lower mean out-degree (solid lines) than *trans* genes, therefore in (D) we perform out-degree conserving shuffles. **(C)** Categories of genes. *cis genes*: genes with a GWS *cis*-eQTL (genes with possible direct effects). *Omnigenes*: genes where including higher orders of propagation increases $R_i^2$ (i.e. QOM $3^{rd}$ > QOM $2^{nd}$ > QOM $1^{st}$ > *cis*). *Higher order QOM* genes: all lower order models explain less than 60% of QOM ∞ expression variance while QOM ∞ explains at least 80% of PRS variance. *PRS* genes: QOM $R_i^2$ is less than 40% of PRS $R_i^2$. **(D)** Out-degree conserving $1^{st}$ and $2^{nd}$ *order* QOM shuffles (distributions), their 90% confidence interval (dashed lines) and models based on "true" network (thick lines). The "true" networks significantly outperform random shuffles. **(E)** Per gene $R_i^2$ and consistency score for examples of all (overlapping) categories of genes from panel (C) (cis genes, omnigenes, PRS genes etc). For clarity, examples with highest $R_i^2$ and consistency> 0.2 are shown for clarity (mean consistency score 0.21). Predicted expression of `YOL116W` is highlighted in panel (F), clade-specific expression of `YKL109W` and `YDR520C` (purple) shown in Fig S10I. **(F)** Per-gene variance explained, $R_i^2$, as a function of variance explained by the PRS model. Errorbars are standard deviations across samples (SI Appendix Sec. 1). Marginal distributions of $R_i^2$ are shown on the sides, shaded area includes genes with above median (13%) PRS. `YOL116W` is highlighted in black.
